# PARP1 Suppression Drives ROS Resistance in Aneuploid Cancer Cells

**DOI:** 10.1101/2025.08.13.670212

**Authors:** Pan Cheng, Angela Mermerian-Baghdassarian, Yufeng Wang, Ze Chen, Helberth M. Quysbertf, Pradeep Singh Cheema, Joseph C. Mays, Xin Zhao, Lizabeth Katsnelson, Sally Mei, Rohini Shrivastava, Mirna Bulatovic, Jiehui Deng, Markus Schober, Kwok-Kin Wong, Teresa Davoli

## Abstract

Aneuploidy—defined as gains and losses of chromosomes—is frequently observed in cancer and has been implicated in promoting tumor progression and metastasis. However, the molecular mechanisms underlying this phenomenon remain poorly understood. By generating new models of aneuploidy, we found that aneuploidy confers remarkable resistance to reactive oxygen species (ROS)-mediated cell death. This resistance is a general consequence of aneuploidy, independent of the specific chromosomes gained or lost. Mechanistically, Poly(ADP-Ribose) Polymerase 1 (PARP1) is suppressed in aneuploid cells, which inhibits PARP1-mediated cell death after ROS (parthanatos). We validated aneuploidy-associated PARP1 suppression across 15 cell models and human tumors, with pronounced effects in metastatic tumors. Importantly, decreased PARP1 levels promote tumor metastasis while increased PARP1 suppresses it. Through a genome-wide CRISPR screen and functional validation, we identified the transcription factor CCAAT/enhancer-binding protein beta (CEBPB) as a critical mediator of PARP1 downregulation and ROS resistance in aneuploid cells. Furthermore, we found that lysosomal dysfunction serves as the upstream mediator of CEBPB activation in aneuploid cells. We propose that aneuploidy-driven CEBPB activation promotes PARP1 suppression, fostering ROS resistance and cancer progression.

**Highlights:**

- Aneuploidy confers resistance to oxidative stress independent of p53 status, karyotype and cell lineage through inhibition of PARP1 expression and activity
- Suppressed PARP1 enhances metastatic potential, while PARP1 restoration suppresses metastatic spread, revealing a novel mechanism linking aneuploidy to cancer progression.
- PARP1 suppression compromises DNA damage repair and cell death to multiple genotoxic stressors, including reactive oxygen species, alkylating agents, and UV radiation.
- A genome-wide CRISPR screen and a CRISPRa screen identifies CEBPB as a critical transcription factor mediating PARP1 regulation and ROS resistance.
- Nuclear CEBPB increases significantly after aneuploidization in experimental systems and in primary human cancers.

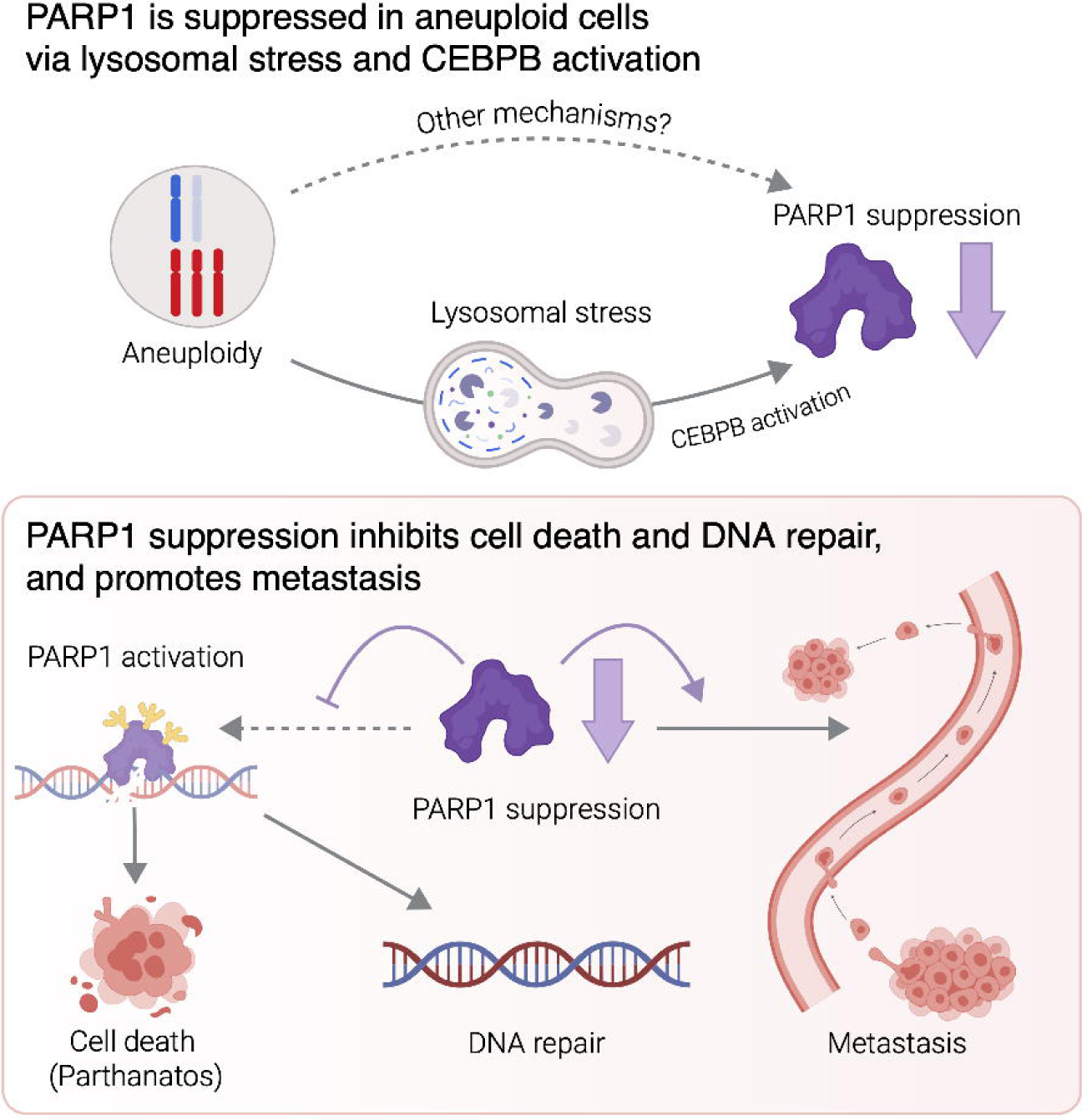

## Introduction

Aneuploidy, a hallmark of cancer^1^, correlates with tumor progression, poor survival, and therapeutic resistance^2–4^. The role of aneuploidy in cancer, however, remains incompletely understood. One prevailing model proposes that aneuploidy promotes tumor-associated phenotypes by altering the dosage of oncogenes and tumor suppressors located on aneuploid chromosomes (**chromosome-specific effects**)^5–7^. The role of the aneuploid state itself (**general effects**) in promoting tumorigenesis, independent of specific chromosomal changes, is more controversial. Aneuploid cells show fitness disadvantages likely due to defects in DNA replication and proteostasis^8,9^, yet resist certain chemotherapies through increased genomic instability and karyotype heterogeneity or through other mechanisms^10,11^. Whether certain general effects of aneuploidy can promote other cancer-associated phenotypes remains unclear^12–14^. A better understanding of how aneuploidy impacts tumor formation and progression could contribute to the development of effective therapies and response biomarkers.

ROS are a double-edged sword in cancer. Low-level ROS contributes to tumorigenesis and cancer development while excessive ROS during tumor progression or anti-cancer therapy can trigger cell death^15,16^. It has been recently shown that oxidative stress in circulating cancer cells inhibits metastasis^17–19^. Therefore, cancer cells tend to enhance their ROS buffering capacity to reduce cell damage, or decrease their sensitivity to ROS-induced cell death, particularly during advanced tumor progression^17^. Interestingly, aneuploidy levels increase with tumor progression and metastasis^2,20^. However, whether a functional relationship between aneuploidy and ROS sensitivity exists and its implications for cancer progression has not been investigated.

Oxidative stress primarily induces DNA single-strand breaks (SSBs), resulting from oxidized phosphate-deoxyribose backbone fragmentation or base excision repair (BER) of oxidized bases^21^. PARP1 rapidly senses SSBs, transfers ADP-ribosyl moieties from nicotinamide-adenine-dinucleotide (NAD+) to various proteins, and facilitates the accumulation and stabilization of repair components at SSB sites^22^. Beyond DNA repair and genome integrity maintenance, PARP1 mediates a ROS-induced cell death mechanism termed parthanatos^23^. In this process, excessive oxidative DNA damage hyperactivates PARP1, depleting cells of the essential NAD+ and adenosine triphosphate (ATP)^24^ and triggering nuclear translocation of mitochondrial apoptosis-inducing factor (AIF)^25^, ultimately leading to cell death. Parthanatos is critical in pathophysiological processes involving cellular stress and excessive DNA damage^23,26^. For example, during cerebral ischemia, PARP1-mediated parthanatos causes neuronal cell death, which PARP1 inhibition can mitigate^27^. However, the role of parthanatos in cancer is much less well understood^28^.

By studying ROS resistance in the context of cancer, we observed PARP1 suppression in aneuploid cells, which enhances resistance to ROS-induced cell death while delaying DNA damage repair. This creates a ‘perfect storm’ benefiting cancer cells where death resistance is coupled with compromised DNA repair. PARP1 modulation affects metastatic potential in both diploid and aneuploid cancer cells. Finally, we identified lysosomal dysfunction, a common feature of aneuploidy^29,30^, activates transcription factor CEBPB, consequently suppressing PARP1 and conferring survival advantages to aneuploid cells under oxidative stress.

## Results

### General aneuploidy confers oxidative stress resistance in acute and chronic models of aneuploidy

To screen potential pro-tumorigenic effects of general aneuploidy (independent of specific chromosome alterations), we developed isogenic aneuploidy models. In a model of **acute aneuploidy** (**Fig. 1A**), immortalized human colon epithelial cells (hCECs) with wild-type (*TP53*WT) or knockout *TP53* (*TP53*KO) were treated with *MPS1/TTK* inhibitors (reversine or AZ3146; MPS1i) for 24-30 hours to induce chromosome missegregation^29^. Aneuploidy was confirmed by altered DNA content (**Fig. 1B**). Karyotypes inferred from single-cell RNA sequencing (see **Methods**) demonstrated that all chromosomes in MPS1i-treated *TP53*KO hCECs were gained or lost with similar frequency (25% of MPS1i-treated cells for each chromosome) (**Fig. S1A**, **Table 1A**).

**Figure 1.**
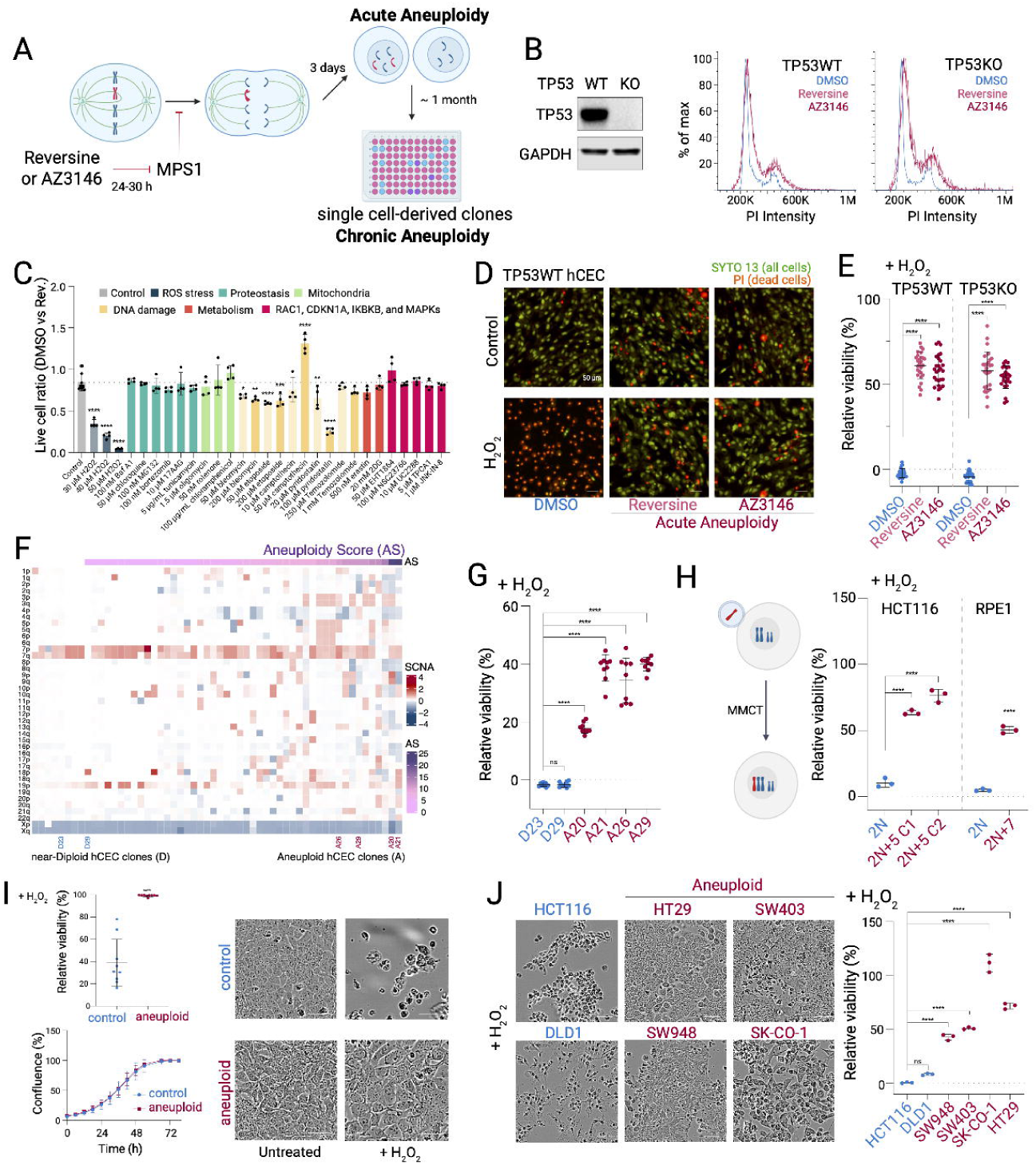
Aneuploidy renders cells resistant to oxidative stress. (**A**) Acute and chronic aneuploidy models. (**B**) DNA content in diploid and aneuploid hCECs. (**C**) Viability of treated diploid and aneuploid hCECs (*TP53*WT). (**D**) Representative images of H_2_O_2_-treated cells (75 μM, 24h). (**E**) Viability quantification from (D). (**F**) Arm-level somatic copy number alterations in hCEC clones (*TP53*KO). (**G**) Viability of H_2_O_2_-treated hCEC clones (50 μM H_2_O_2_ for 1h + 24h release). (**H**) Viability of H_2_O_2_-treated HCT116 or RPE1 clones (4mM and 100 μM, respectively, 24h). (**I**) Viability (top left) and representative images (right) of H_2_O_2_-treated KP cells (2 mM, 24h). Proliferation curves (bottom left). (**J**) Representative images and viability of H_2_O_2_-treated colon cancer cells (4 mM, 24 h). One-way ANOVA with Dunnett’s test (C, D, HCT116 in H). Tukey’s test (G, J). unpaired t-test (RPE1 in H, I). Scale bar, 50 μm.

We next compared resistance to various cellular stresses between diploid and aneuploid *TP53*WT hCECs (DMSO versus MPS1i) (**Fig. 1C**, **Fig. S1B**). GFP-labeled diploid and unlabeled aneuploid hCECs were treated with drugs inducing various cellular stresses, including oxidative stress, proteostasis stress, energy stress, DNA damage, and metabolic stress. Inhibitors of RAC1, CDKN1A, IKBKB, and MAPKs were also included (**Fig. 1C, Fig. S1B, Table 1B**). Remarkably, compared to diploid ones, aneuploid cells were much more resistant to oxidative stress caused by hydrogen peroxide (H_2_O_2_) in a concentration-dependent manner (**Fig. 1C, Fig. S1B).** Importantly, H_2_O_2_ resistance was independent of *TP53* status (**Fig. 1D-E**).

To assess MPS1i off-target effects, hCECs were treated with MPS1i (or DMSO) at high confluency, where cell division and aneuploidy formation are suppressed but potential off-target effects remain (**Fig. S1C**). These MPS1i-treated but non-aneuploid hCECs exhibited increased ROS-induced cell death compared to controls (30-50% vs 14%) (**Fig. S1C**). This observation, along with similar results from different MPS1i, suggests that ROS resistance in MPS1i-treated cells depends on aneuploidy. Next, we validated this finding using other cells, including non-cancerous human cells from different tissues (**Fig. S1D-E**). Furthermore, we explored the impact of the cell cycle state on oxidative stress resistance. Arresting hCECs at G1 or G1/S phase (using palbociclib or thymidine) increased ROS resistance in both diploid and aneuploid cells by 20-40%. However, arrested aneuploid cells maintained significantly higher resistance, with ∼30% greater relative viability compared to diploid cells (**Fig. S1F**).

To examine aneuploidy effects on ROS resistance over a longer timeframe, we generated a **chronic aneuploidy** model^31^. Using reversine-treated *TP53*KO hCECs, we generated 55 single-cell-derived isogenic hCEC clones with varying aneuploidy levels, assessed by whole-genome sequencing (**Fig. 1A, 1F**, **Table 1C**). Two representative near-diploid and four aneuploid clones (≥5 gains or losses of chromosomes or chromosome arms, with stable karyotypes) were compared for their oxidative stress resistance (**Fig. 1F**, **Fig. S1G**). The aneuploid clones showed enhanced resistance compared to near-diploid clones (**Fig. 1G, Fig. S1H-K**). Increased resistance to oxidative stress was also confirmed using aneuploid clones of HCT116 (a human colon cancer cell line) and RPE1^32^ (**Fig. 1H**).

We further validated these findings by inducing aneuploidy in near-diploid mouse KP (*Kras^G12D^* and *Trp53*^-/-^) lung adenocarcinoma model^33,34^ using an adapted KaryoCreate strategy (**Fig. 1I, S1L**). Aneuploid KP cells displayed similar proliferation but ∼60% higher resistance to oxidative stress compared to near-diploid controls (**Fig. 1I**), further supporting that ROS resistance in aneuploid cells is not caused by cell cycle alterations. Finally, in a panel of colon cancer cell lines, high-aneuploidy lines (HT29, SW403, SW948, and SK-CO-1) (**Table 1D**) were more resistant to oxidative stress compared to near-diploid ones (HCT116 and DLD1) with (40-110% higher viability, **Fig. 1J**, **Fig. S1M**).

We also confirmed aneuploidy-induced ROS resistance using alternative ROS inducers, supplementing the culture medium with xanthine oxidase and xanthine to produce both O_2_^-^ (superoxide) and H_2_O_2_^35^ (**Fig. S1N**). Collectively, our data across multiple aneuploidy models provide robust evidence that aneuploidy renders cells resistant to oxidative stress, regardless of *TP53* status, cell cycle, altered chromosomes, or cell lineage.

### Aneuploid cells resist ROS despite antioxidant levels comparable to diploid cells

The aneuploidy-induced ROS resistance could result from heightened antioxidant defenses. NRF2 (nuclear factor erythroid 2-related factor 2) is the first-line defense against oxidative stress^17^, translocating to the nucleus upon activation to initiate antioxidant gene transcription^36^. We examined *NRF2* activation and found that acute aneuploid hCECs showed no significant increase in NRF2 nuclear translocation (**Fig. S2A**). Aneuploid hCEC clones even displayed slightly reduced NRF2 nuclear translocation versus near-diploid clones. Consistently, *NRF2* knockout did not abolish the survival advantage of aneuploid hCECs relative to diploid controls under oxidative stress (**Fig. S2B**). Furthermore, knockdown of key antioxidant genes, *GPX1* and *CAT*, which are crucial for direct cellular H_2_O_2_ elimination, did not abolish ROS resistance in aneuploid cells (**Fig. S2C**). Collectively, these findings suggest that the increased oxidative stress resistance in aneuploid cells is not mediated by enhancement of cellular antioxidant defenses via *NRF2*, *GPX1,* or *CAT*.

We then compared ROS-induced cellular damage between diploid and aneuploid hCECs (**Fig. 2A**, **Fig. S2D-E**). Alkaline comet assay revealed that H_2_O_2_ exposure induced a level of DNA damage (double and single-strand breaks) that is comparable or slightly higher in aneuploid versus diploid hCECs **(Fig. 2A, Fig. S2D**). This result was corroborated by the similar levels of the DNA damage marker phosphorylated histone H2AX (Ser139; γH2AX) between diploid and aneuploid cells after H_2_O_2_ (see **Fig. 3M**, **Fig. S3H-K**). Furthermore, H_2_O_2_ exposure resulted in similar lipid peroxidation levels in diploid and aneuploid hCECs (**Fig. S2E**). These findings indicate that the resistance of aneuploid cells to oxidative stress is not due to enhanced antioxidant defenses, as no reduction in oxidative DNA or lipid damage was observed in aneuploid compared to diploid cells following H_2_O_2_ exposure.

**Figure 2.**
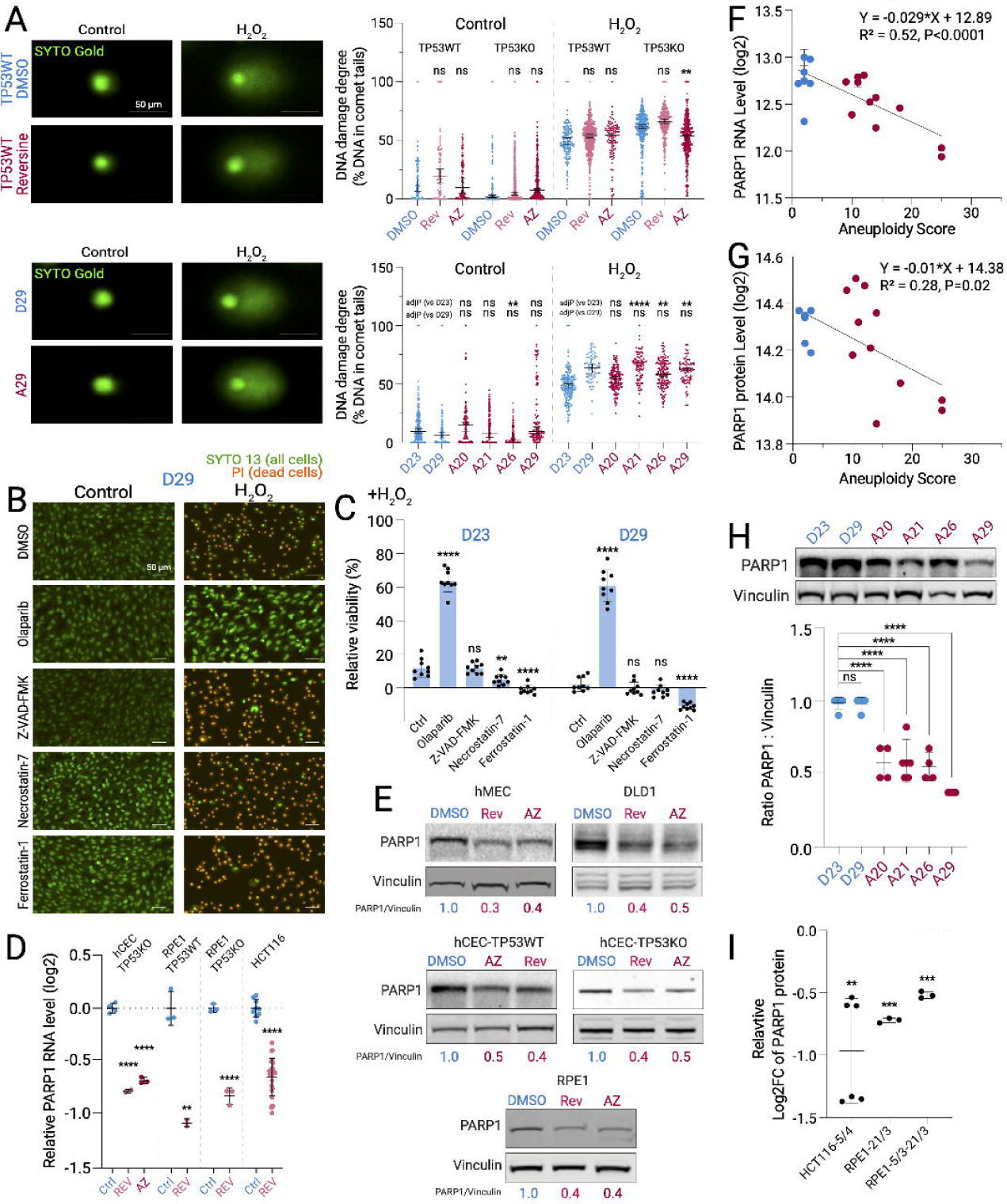
PARP1 is suppressed in aneuploid cells and required for ROS-induced cell death. (**A**) DNA damage in diploid and aneuploid hCECs after H_2_O_2_ (50 μM, 1h). Left: representative images. Right: quantification of damaged DNA. Medians with 95% CI. (**B**) Representative images of H_2_O_2_-treated hCECs (50 μM, 1h) with cell death inhibitors (2h pretreatment + 24h post-treatment). (**C**) Viability quantification from (B). (**D**) Relative PARP1 RNA levels (Log2) in acute aneuploidy. (**E**) PARP1 protein levels in acute aneuploidy. (**F-G**) Correlation between aneuploidy degree and PARP1 RNA (F) or protein (G) levels in hCEC clones. (**H**) PARP1 protein levels in hCEC clones. (**I**) Relative PARP1 protein levels in HCT116 and RPE1 clones (data from Stingele et al., 2012). Notation: cell line-chromosome-copy number Kruskal-Wallis one-way ANOVA with Dunnett’s test (A). One-way ANOVA with Dunnett’s test (C, hCEC in D); Tukey’s test (H). Unpaired t-test (RPE1, *TP53*KO RPE1, and HCT116 in D). One sample t-test against zero (I). Scale bar, 50 μm.

**Figure 3.**
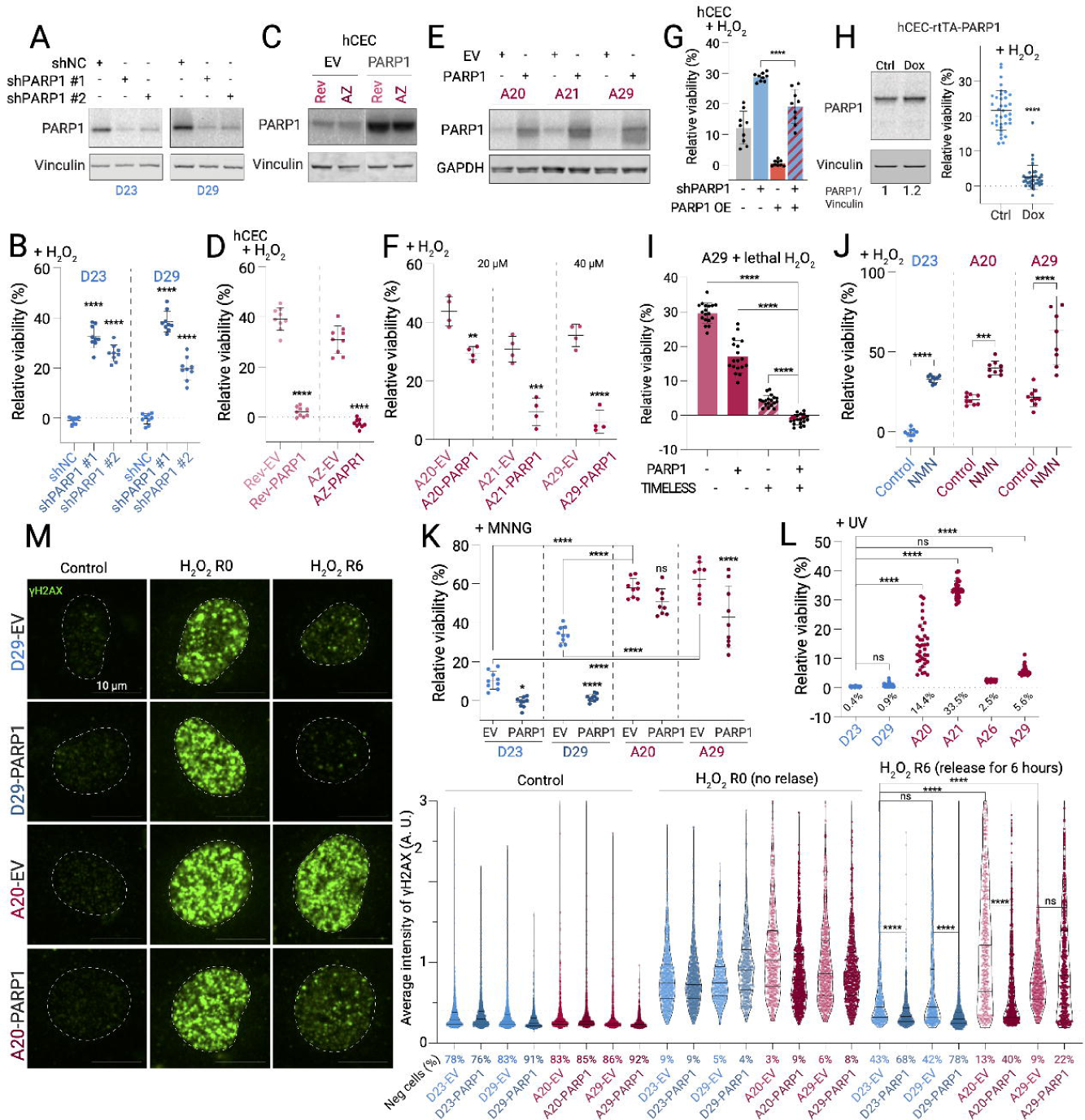
Cell death and DNA repair mediated by PARP1 are suppressed in aneuploid cells by PARP1 downregulation. (**A**) PARP1 levels in near-diploid hCEC clones with PARP1 knockdown. (**B**) Viability of cells from (A) after H_2_O_2_. Mean ± SD of 9 fields (**C**) PARP1 levels in the acute aneuploidy (*TP53*WT hCEC) with PARP1 overexpression. Dashed lines indicate spliced lanes. (**D**) Viability of cells from (B) after H_2_O_2_. (**E**) PARP1 levels in aneuploid hCEC clones with PARP1 overexpression. (**F**) Viability of cells from (E) after H_2_O_2_. (**G**) Viability after H_2_O_2_ treatment with *PARP1* knockdown, PARP1 overexpression, or both. (**H**) PARP1 levels (left) and H₂O₂-induced cell viability (right) in hCECs with doxycycline-inducible PARP1 overexpression. (**I**) Viability after H_2_O_2_ (40 μM) with PARP1or TIMELESS overexpression or both. (**J**) Viability after H_2_O_2_ in hCEC clones with NMN pretreatment (180 μM, 24h). (**K**) Viability after MNNG treatment (62.5 μM, 24 hours) in hCEC clones with control or PARP1 overexpression. (**L**) Viability after UV (50 J/m^2^ + 5-day release) in hCEC clones. Average viability indicated below dashed line. (**M**) γH2AX before and after H_2_O_2_ (50 μM, 1h) in hCEC clones ± PARP1 overexpression. Left: images (scale bar, 10 μm). Right: violin plots showing median and quartiles (>460 cells/condition). R0/R6: 0 or 6h post-H_2_O_2_. All p-values in Table 3. γH2AX-negative threshold set at ∼80% for untreated clones; percentages shown below X-axis. One-way ANOVA with Dunnett’s test (B); with Tukey’s test (G, I, J, K, L). Unpaired t-test (D, F, H). Kruskal-Wallis one-way ANOVA with Dunnett’s test (M).

### PARP1, a key mediator of ROS-induced cell death, is suppressed in aneuploid cells

Since no reduction in oxidative DNA or lipid damage was observed in aneuploid cells, we investigated whether aneuploid cells better resist ROS-induced cell death despite comparable levels of cellular damage. To determine the specific cell death pathway activated by oxidative stress, we pretreated hCECs with inhibitors of apoptosis, necrosis, ferroptosis, and PARP1-mediated parthanatos (olaparib). Strikingly, only olaparib significantly rescued H_2_O_2_-induced cell death (**Fig. 2B-C, Fig. S2F**). This indicates that elevated oxidative stress triggers parthanatos in our cell model.

PARP1, the molecular target of olaparib, is crucial for parthanatos: it senses DNA damage caused by ROS or other stimuli, and in the presence of high levels of damage, it initiates parthanatos by exhausting NAD+ and ATP or by triggering AIF nuclear translocation^24,25^. Importantly, we observed a consistent decrease in PARP1 RNA (∼40-50%) and protein (∼50-60%) levels in aneuploid cells compared to diploid controls across multiple cell line models (**Fig. 2D-E**, **Table 2A-B**), independent of their transformation state, *TP53* status, or cell lineage. PARP1 downregulation was also observed in previous reports on reversine-treated RPE1 cells^29^, glioma cells^37^, and mouse embryos^38^, while none of these studies focused on PARP1 or parthanatos. Furthermore, we probed for PARP1 activity and observed reduced poly(ADP-ribose) (pADPr) synthesis in MPS1i-treated aneuploid cells after H_2_O_2_ exposure (**Fig. S2G**). Importantly, the decline in PARP1 protein cannot be attributed to caspase-mediated cleavage^39^, as it persisted with a pan-caspase inhibitor (**Fig. S2H**). It is important to highlight that *PARP1* is haploinsufficient^40^, thus a partial change in gene dosage can profoundly impact cell state and function.

The negative correlation between aneuploidy degree and PARP1 expression at RNA and protein levels was also validated in the isogenic aneuploid and diploid hCEC clones (RNA: Spearman R = -0.58, p = 0.01; protein: Spearman R = -0.49, p = 0.05; **Fig. 2F-H**, **Table 2C-D**). HCT116 clones and RPE1 clones (**Fig. 2I**, **Fig. S2I-J**), each carrying different aneuploidies, exhibited lower PARP1, suggesting that PARP1 downregulation is independent of specific altered chromosomes. Finally, aneuploidy-associated PARP1 downregulation was also validated using KP lung cancer cells (**Fig. S2K**) and human colon cancer cell lines (see **Fig. 4A**). Taken together, these data suggest that PARP1 downregulation is a common feature of aneuploid cells, regardless of *TP53* status, altered chromosomes, cell lineage, or transformation state.

**Figure 4.**
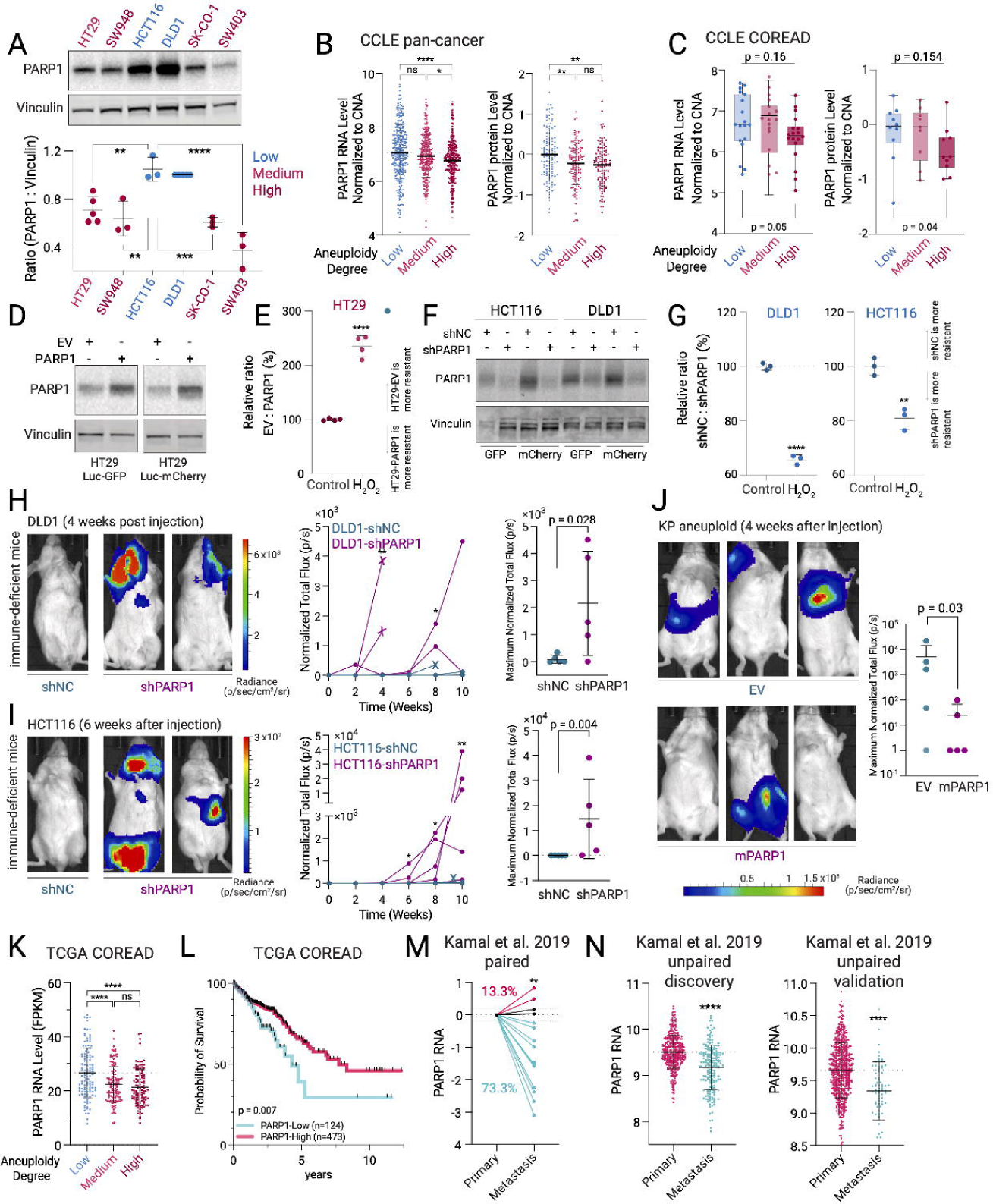
PARP1 suppression promotes metastasis of aneuploid cancer cells. (**A**) PARP1 levels in indicated cell lines. (**B-C**) SCNA-normalized *PARP1* RNA or protein levels in pan-cancer (B) or in colon cancer cell lines (C) with different aneuploidy degrees. (**D**) PARP1 protein in HT29 cells ± *PARP1* overexpression. (**E**) Relative viability of HT29-mCherry-EV to HT29-GFP-PARP1 when co-cultured ± H_2_O_2_ (2 mM, 24h). (**F**) PARP1 protein in HCT116 and DLD1 cells ± *PARP1* knockdown. (**G**) Relative viability of control vs PARP1-knockdown cells when co-cultured ± H_2_O_2_ (2 mM, 24h). (**H-I**) Bioluminescence images (BLI) of NSG mice with DLD1 or HCT116 ± *PARP1* knockdown. Left: representative images. Middle: total flux over time. Right: maximum values. Mean ± SD, n=5 mice/group. Crosses indicate endpoints. (**J**) BLI of immune-intact mice with aneuploid mouse KP lung cancer cells ± *PARP1* overexpression. Left: representative images. Right: maximum total flux signal. Mean + SD, n=5 mice/group. (**K**) *PARP1* RNA levels in TCGA COREAD patients without PARP1 SCNA. (**L**) Kaplan-Meier survival of TCGA COREAD patients stratified by PARP1 expression. (**M**) *PARP1* RNA levels in primary COREAD tumors and their paired metastases (n=15 patients, GSE131418). (**N**) *PARP1* RNA in unpaired primary tumors versus metastases of COREAD patients (discovery cohort: 333 primary, 184 metastases; validation cohort: 545 primary, 72 metastases, GSE131418). One-way ANOVA with Tukey’s test (A, B, K). Kruskal-Wallis test, followed by one-tailed t-test comparing low vs high aneuploidy groups (C). Unpaired t-test (E, G, N). Two-tailed Mann-Whitney test (H-I). One-tailed Mann-Whitney test (J). Log-rank (Mantel-Cox) test (L). One Sample Wilcoxon signed-rank test (M).

### PARP1 suppression reduces aneuploid cell sensitivity to PARP1-mediated cell death induced by ROS, alkylating agents, or UV radiation

While *PARP1* is considered a pro-survival gene that senses DNA damage and promotes DNA repair^22^, its hyperactivation can trigger parthanatos, the most common form of regulated necrosis induced by DNA damage^26,41^. Given our findings of *PARP1* suppression in aneuploid cells (**Fig. 2**), we hypothesized that this reduction contributes to their resistance to ROS-induced (or more generally, PARP1-mediated) cell death. To test this hypothesis, we modulated *PARP1* expression through knockdown/knockout or overexpression and assessed corresponding changes in cell sensitivity to H_2_O_2_, an activator of PARP1-mediated cell death.

shRNA- or CRISPR-mediated *PARP1* inhibition conferred H_2_O_2_ resistance to near-diploid hCEC clones, compared to control cells (**Fig. 3A-B**, **Fig. S3E-F**). Conversely, PARP1 overexpression decreased H_2_O_2_ resistance in both acutely and chronically aneuploid hCECs, reducing relative viability compared to empty vector controls (**Fig. 3C-F, Fig. S3A**). Notably, the resistance induced by the shRNA targeting *PARP1* 3’-UTR was reversed by overexpressing shRNA-resistant *PARP1* cDNA (**Fig. 3G**). Consistent with observations in hCECs, *PARP1* overexpression in the high-aneuploidy colon cancer cell line (HT29) reduced H_2_O_2_ resistance (see **Fig. 4D-E**, **Fig. S4A**), while *PARP1* knockdown in the near-diploid colon cancer cell lines (HCT116 and DLD1) enhanced resistance (see **Fig. 4F-G**, **Fig. S4B**). Notably, using doxycycline-inducible PARP1 expression, we found that a 20% increase in PARP1 expression sufficiently reduced H_2_O_2_ resistance, decreasing relative viability by ∼20% (**Fig. 3H**).

To examine additional factors that may work with PARP1 to contribute to aneuploidy-associated oxidative stress resistance, we next investigated PARP1-associated proteins that have been shown to promote its function. One of the most prominent is Timeless Circadian Regulator (*TIMELESS*), which physically interacts with PARP1 and promotes its recruitment to DNA damage foci and subsequent activation^42^. *TIMELESS* expression was consistently decreased at RNA levels in acutely aneuploid cells compared to diploid controls (**Fig. S3B**) and inversely correlated with aneuploidy degree at RNA and protein levels in the chronic aneuploidy model (**Fig. S3C**). We found that in the highly-resistant aneuploid clone A29, co-overexpression of *PARP1* and *TIMELESS* reduced H_2_O_2_ resistance more than overexpression of either gene alone (**Fig. 3I**, **Fig. S3D**), suggesting TIMELESS downregulation may enhance ROS resistance induced by *PARP1* suppression in aneuploid cells. Given the strong effect of TIMELESS overexpression on ROS sensitivity, we investigated the necessity of *PARP1* downregulation in *TIMELESS*-mediated ROS sensitivity. We knocked out *PARP1* in A29 cells overexpressing *TIMELESS*. Strikingly, *PARP1* knockout completely abolished the sensitizing effect of *TIMELESS* overexpression on H_2_O_2_-induced cell death (**Fig. S3L**), demonstrating that *PARP1* is the primary effector of oxidative stress sensitivity and that *TIMELESS* acts through *PARP1* to modulate this response. While *PARP1* catalyzes 85-95% of total cellular pADPr in response to DNA damage, *PARP2* has a similar function and is also targeted by olaparib^43^. However, *PARP2* knockout did not protect hCECs from H_2_O_2_ (P = 0.22; **Fig. S3E**), arguing against its involvement in aneuploidy resistance to parthanatos. Macrophage migration inhibitory factor (*MIF*), a late-stage parthanatos nuclease, cleaves genomic DNA^44^. *MIF* knockout partially protected hCECs from H_2_O_2_, but less effectively than *PARP1* knockout (relative viability increased by 20-38% for *PARP1* knockout vs 7-14% for *MIF* knockout, **Fig. S3F**).

Furthermore, we investigated whether mitigating NAD+ depletion, a typical consequence of PARP1 hyperactivation^23,45^, inhibits cell death after H_2_O_2_ treatment. Supplementation with nicotinamide mononucleotide (NMN, a NAD+ precursor) increased the viability of H_2_O_2_-treated hCEC clones (**Fig. 3J**). Finally, we applied agents other than H_2_O_2_-induced ROS to induce DNA damage and PARP1 activation: N-methyl-N′-nitro-N-nitrosoguanidine (MNNG), a common alkylating agent, and UV radiation, both known to trigger PARP1-mediated cell death^41^. Remarkably, aneuploid hCEC clones were more resistant to these stimuli than near-diploid clones in a PARP1-dependent manner (**Fig. 3K-L**). Collectively, these results suggest that PARP1 downregulation confers resistance to PARP1-mediated cell death in aneuploid cells, whether induced by ROS, alkylating agents, or UV radiation.

### PARP1-mediated DNA damage repair is impaired in aneuploid cells

Given the crucial role of *PARP1* in this process^22^ and its downregulation in aneuploid cells, we investigated the impact of *PARP1* suppression on DNA damage repair in aneuploid cells. We transiently exposed isogenic near-diploid and aneuploid hCEC clones to genotoxic insults and collected cells before (control) and after stimulation (with no observed cell death prior to collection). DNA damage was assessed by measuring γH2AX^46^ (**Fig. S3G**). Seven genotoxic agents inducing DNA damage through distinct mechanisms were tested: H_2_O_2_ (oxidative damage), MNNG (base alkylation), camptothecin (topoisomerase I inhibition), etoposide (topoisomerase II inhibition), pyridostatin (C-quadruplex stabilization), doxorubicin (DNA intercalation), and bleomycin (DNA synthesis inhibition). We found that repair of H_2_O_2_- and MNNG-induced DNA damage was significantly slower in aneuploid clones compared to diploid cells (see **Methods**), whereas repair of DNA damage induced by other genotoxic agents was not (**Fig. S3H-I**). After six hours of recovery from H_2_O_2_ treatment, median γH2AX intensity decreased by ∼40% in near-diploid clones but by only 15% (A29) or increased by 20% (A20) in aneuploid clones (**Fig. S3H-I**). Similarly, MNNG-induced DNA damage repair was less efficient in aneuploid clones, with median γH2AX intensity increasing by 54-103% compared to -16% to 19% in near-diploid clones after six hours of recovery (**Fig. S3H-I**). This likely reflects the strong reliance on PARP1 for repairing H_2_O_2_- and MNNG-induced DNA damage, whereas other agents engage multiple, less PARP1-dependent repair pathways^46^ (see **Discussion**). The suppressive effect of PARP1 downregulation on oxidative DNA damage repair was further validated by analyzing PARP1-overexpressing cells. PARP1 overexpression promoted oxidative DNA damage repair in both acute and chronic aneuploidy models (**Fig. 3M, Fig. S3J**, **Table 3**). Importantly, we excluded the possibility that γH2AX dynamics resulted from altered H2AX protein levels, as γH2AX changes were independent of total H2AX level fluctuations (**Fig. S3K**). Additionally, the γH2AX decrease in the near-diploid clones 6 hours after H_2_O_2_ exposure is likely unrelated to programmed cell death, as previous studies showed PARP1-mediated cell death does not cause collateral γH2AX decrease^47^. Taken together, our data indicate that aneuploid cells, besides surviving H_2_O_2_ or MNNG-induced cell death, may be more prone to accumulating DNA damage due to suppressed PARP1-mediated DNA repair. The combination of death resistance and DNA damage accumulation may collectively promote tumorigenesis and cancer evolution (see **Discussion**).

### PARP1 downregulation promotes tumor progression of aneuploid cancer cells

Given the decreased PARP1 expression across different aneuploidy models (**Fig. 2**), we asked whether the same was true in human cancer cell lines and in primary tumors with varying aneuploidy levels. Western blot (WB) analysis of four high-aneuploidy (HT29, SW948, SK-CO-1, SW403) and two near-diploid (HCT116, DLD1) colon cancer cell lines revealed lower PARP1 expression in high-aneuploidy lines (all cell lines have two copies of *PARP1* gene, **Fig. 4A**, **Table 1D**). Analysis of pan-cancer cell lines in the CCLE dataset, evenly grouped into three aneuploidy levels, revealed significant aneuploidy-dependent decreases in PARP1 expression at both SCNA-normalized RNA and protein levels (RNA mean log2FC = -0.3, protein mean log2FC = -0.25, high vs low aneuploidy, p<0.0001, p=0.0013; **Fig. 4B**). This association extended to colorectal cancer cell lines from the CCLE dataset (**Fig. 4C**).

Furthermore, consistent with our findings in hCECs that PARP1 expression inversely correlates with cell resistance to ROS-induced death (**Fig. 3**), *PARP1* overexpression attenuated ROS resistance in high-aneuploidy colon cancer cells (**Fig. 4D-E**, **Fig. S4A**), while *PARP1* knockdown enhanced ROS resistance in near-diploid colon cancer cells (**Fig. 4F-G**, **Fig. S4B**). These results further support our model, indicating that PARP1 expression levels critically determine cancer cell sensitivity to ROS-induced cell death (**Fig. 3**).

Since metastasis requires cancer cells to withstand high oxidative stress during migration and colonization^17–19,48^, we investigated whether PARP1 levels influence dissemination potential of diploid and aneuploid cancer cells. We found *PARP1* suppression enhanced dissemination potential of near-diploid human colon cancer cells in immunodeficient mice (**Fig. 4H-I**). DLD1 and HCT116 cells transduced with *PARP1* shRNA exhibited significantly increased bioluminescence signals four or six weeks post-injection compared to controls with non-targeting shRNA, respectively (n=5 mice per group; p = 0.007 for DLD1 at week 4, p = 0.03 for HCT116 at week 6; **Fig. 4H-I**). Importantly, *PARP1* knockdown did not affect cell proliferation *in vitro* (**Fig. S4C**). Furthermore, we assessed the impact of PARP1 levels on murine aneuploid cancer cells in immune-intact mice. We intravenously injected aneuploid mouse KP lung cancer cells (**Fig. 1I**, **Fig. S1J**) into B6 Albino mice. Consistently, mouse PARP1 overexpression decreased their tumor dissemination without affecting *in vitro* proliferation, with 40% of mice developing tumors in the PARP1 OE group compared to 80% in empty vector controls (**Fig. 4J**, **Fig. S4D**). These data indicate that PARP1 downregulation may promote metastasis of aneuploid cancer cells in both immune-deficient and immune-intact mice.

We next examined PARP1 levels in human primary tumors to determine whether it changes with tumor stage and patient survival. In line with our *in vivo* findings, among TCGA colorectal adenocarcinoma (COREAD) patients lacking PARP1 SCNA^49^, PARP1 expression was lower in high-versus low-aneuploidy tumors (**Fig. 4K**, **Table 4**), and lower PARP1 expression was associated with shorter survival (**Fig. 4L**, **Table 4**). This aligns with previous studies showing that late-stage COREAD tumors, typically characterized by higher aneuploidy degrees^50^, exhibit lower PARP1 expression and activity compared to early-stage tumors^51^. Furthermore, we compared PARP1 levels in primary and metastatic COREAD samples and found decreased PARP1 expression in metastases compared to primary tumors in both paired (p < 0.01; **Fig. 4M**) and unpaired sample cohorts^52^ (**Fig. 4N**, **Fig. S4E**). Altogether, these findings support a role for *PARP1* downregulation in promoting metastasis.

### CRISPR screen identifies lysosomal stress is a potential mediator of PARP1 downregulation in aneuploid cells

Next, we investigated the mechanisms of *PARP1* regulation, specifically its downregulation in aneuploid cells. We conducted fluorescence-activated cell sorting (FACS)-based genome-wide CRISPR screens to identify genes modulating PARP1 expression, using endogenous PARP1 levels as the readout (**Fig. 5A-B** and **Fig. S5A**).

**Figure 5.**
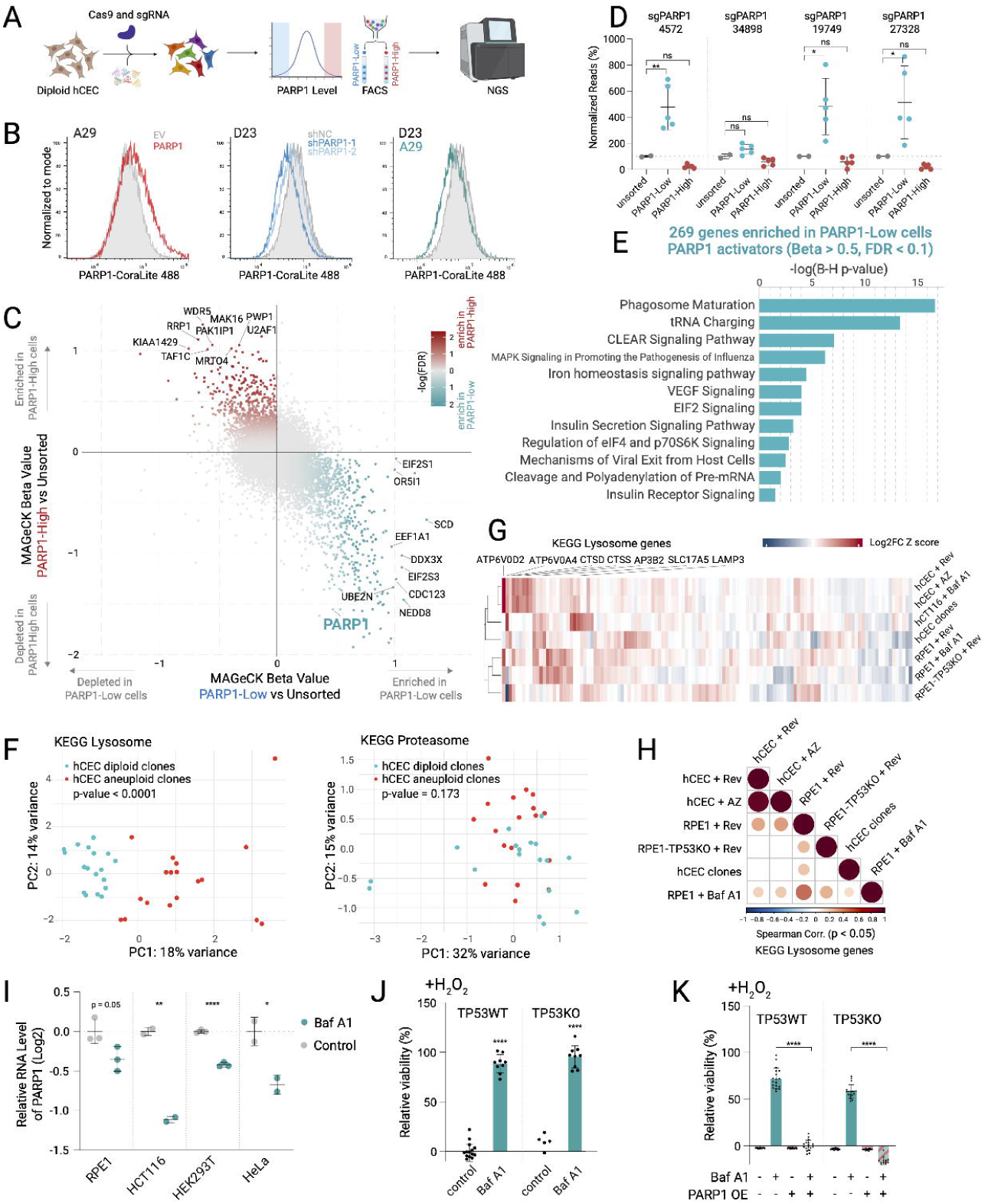
Genome-wide CRISPR screen reveals lysosomal stress as a mediator of PARP1 downregulation in aneuploid cells. (**A**) FACS-based genome-wide CRISPR screen to find PARP1 regulators. (**B**) Validation of PARP1 antibody. (**C**) Enrichment and depletion of sgRNAs compared to unsorted samples. (**D**) Normalized sgPARP1 read counts in sorted vs unsorted samples. (**E**) Pathway analysis of 269 PARP1 activators. (**F**) PCA of lysosomal (left) and proteasomal (right) gene expression in near-diploid (blue) and aneuploid (red) hCEC clones. Group separation assessed by 10,000 permutations using Euclidean distances in PC1-PC2 space. (**G**) Heatmap of log2FC z-scores for KEGG lysosomal genes in aneuploid cells and Bafilomycin A1 (Baf A1)-treated cells. (**H**) Spearman correlation of lysosomal gene expression changes in aneuploid cells and Baf A1-treated RPE1 cells. (**I**) *PARP1* mRNA levels (Log2) in Baf A1-treated cells. (**J**) Viability of hCECs pre-treated with Baf A1 (5 nM, 24h) followed by H_2_O_2_ (200 μM, 24 h). (**K**) Viability after H_2_O_2_ in hCECs ± Baf A1 pre-treatment or PARP1 overexpression. One-way ANOVA with Dunnetts’ test (D); with Tukey’s test (K). Unpaired t-test (I, J).

SgRNAs were sequenced from PARP1-low, PARP1-high, and unsorted cells. We anticipated sgRNAs targeting putative *PARP1* activators to be enriched in PARP1-low cells compared to unsorted cells, while sgRNAs targeting putative *PARP1* suppressors would be enriched in PARP1-high cells. Using MAGeCK to calculate enrichment scores for each gene, we revealed a negative correlation between gene enrichment in PARP1-low and PARP1-high populations (p < 2e-16, rho = -0.22, **Fig. 5C**, **Table 5A**), indicating exclusive enrichment of genes in populations with different PARP1 levels. Importantly, sgRNAs targeting *PARP1* were enriched in the PARP1-low population and depleted from the PARP1-high population (beta = 0.48 and -1.55, p-value = 0.003 and 0 for PARP1-low and PARP1-high populations, respectively; **Fig. 5C-D**), confirming the screen’s success.

The screen identified 269 *PARP1* activators and 155 suppressors (FDR < 0.1, beta > 0.5, **Table 5B**). QIAGEN Ingenuity Pathway Analysis (IPA) revealed that *PARP1* activators were enriched in pathways related to phagosome maturation (p-value = 1.7e-17), tRNA charging (p-value = 3.7e-14) among others. *PARP1* suppressors were predominantly associated with RNA polymerase I complex assembly (p-value = 1.2e-6) (**Fig. 5E**, **Fig. S5B**, **Table 5C-D**).

To identify genes and pathways that may mediate PARP1 downregulation in aneuploid cells, we examined how the expression of these *PARP1* activators and suppressors changes in aneuploid compared to diploid cells (**Fig. S5C**). Aneuploidy-associated *PARP1* regulators were identified by finding *PARP1* activators that are downregulated in aneuploid cells and *PARP1* suppressors that are upregulated in aneuploid cells (**Fig. S5D, Table 5E**). Phagosome maturation pathway (p-value = 6.4e-03), consisting of many lysosome genes, was enriched in these aneuploidy-associated PARP1 regulators (**Fig. S5E**, **Table 5F**), suggesting that lysosomal dysregulation in aneuploid cells may cause PARP1 downregulation.

Lysosomal stress is characteristic of aneuploidy^29^, with aneuploid cells showing transcriptional profiles similar to cells treated with the lysosomal V-ATPase inhibitor bafilomycin A1 (Baf A1)^30^. Our screen showed that knockout of lysosomal genes resulted in PARP1 downregulation (phagosome maturation pathway in **Fig. 5E** and **Fig. S5E**, individual lysosomal *V-ATPase* genes in **Fig. S5F**). Thus, we hypothesized that aneuploidy-associated lysosomal stress induces PARP1 downregulation and tested this hypothesis. First, principal component analysis (PCA) of RNAseq data from 8 near-diploid and 11 aneuploid hCEC clones revealed that lysosomal, but not proteasomal, genes differentiated near-diploid from aneuploid cells (p-value < 0.0001 and = 0.173, respectively, **Fig. 5F**), indicating a distinct lysosomal gene expression profile in aneuploid cells. Second, expression changes of these lysosomal genes and the whole transcriptome in aneuploid cells correlated with those in cells treated with the lysosome inhibitor Baf A1 (Spearman p-value < 0.05, **Fig. 5G-H**, **Fig. S5G**), suggesting that aneuploid cells experience Baf A1-like lysosomal stress. Third, analysis of four published datasets^29,53–55^ revealed that lysosome inhibitor Baf A1 treatment reduced PARP1 expression in various cell lines (Log2FC: -0.350 to -1.112, **Fig. 5I**). Reduction in PARP1 after Baf A1 treatment was further validated by WB analysis in our hCECs (**Fig. S5H**). Conversely, proteasome inhibition^29,56–58^ did not affect PARP1 expression (p-value > 0.05, **Fig. S5I**), highlighting the specificity of lysosome inhibition for PARP1 downregulation. Finally, Baf A1 treatment protected hCECs (both *TP53*WT and *TP53*KO) from H_2_O_2_-induced cell death (91-107% increase in relative viability, **Fig. 5J**). This lysosome dysfunction-induced resistance to oxidative stress was eliminated by PARP1 overexpression (p-value < 0.0001, **Fig. 5K**). These results suggest that lysosomal stress promotes PARP1 downregulation and corresponding oxidative stress resistance in aneuploid cells.

### CEBPB is an aneuploidy-induced transcription factor that suppresses PARP1 expression

Decreased PARP1 RNA levels in cells with aneuploidy or lysosomal dysfunction (**Fig. 2**, **Fig. 5**) suggested that these conditions may alter *PARP1* transcription. We therefore sought to identify transcription factors (TFs) that may be responsible for *PARP1* transcriptional dysregulation in these settings. We integrated our PARP1 CRISPR screen (**Fig. 5C**) with TF databases^59–61^ to identify TFs influencing *PARP1* expression (**Fig. 6A**). Additionally, we examined expression changes of these TFs in aneuploid and lysosome-dysfunction cells, searching for downregulation of TFs promoting *PARP1* expression and upregulation of those suppressing *PARP1* expression (**Fig. 6A**, see Methods). These analyses identified CEBPB as a potential suppressive TF of *PARP1*, with consistently elevated expression in various aneuploid and lysosome-dysfunction cell models (p<0.05, **Fig. 6B-F**). CEBPB ChIP-Seq^62^ confirmed its binding near the *PARP1* transcription start site upon activation by dexamethasone (*CEBPB* activator) (reproducible peaks identified by the Irreproducibility Discovery Rate algorithm, **Fig. 6B**). IF analysis of hCECs showed a significant increase in nuclear CEBPB signal in aneuploid cells compared to near-diploid cells in both the chronic and acute aneuploidy models (**Fig. 6D**). CEBPB upregulation was also confirmed in our transcriptomic and proteomic datasets in both acute (**Fig. 6C**) and chronic models (**Fig. 6E**). Lysosome dysfunction induced by Baf A1 also increased CEBPB levels across multiple lines (**Fig. 6F**) and enhanced nuclear localization in hCECs with *TP53*WT or *TP53*KO (**Fig. S6A**). *CEBPB* activation was also supported by transcription factor enrichment analysis using ChEA3 (ENCODE dataset)^63^, showing significant overlap between CEBPB target genes and highly upregulated genes (log2FC > 1 & FDR < 0.01) in reversine-treated hCECs (ranked 3rd, FDR = 6.38E-5) and Baf A1-treated RPE1 cells (ranked 5th, FDR = 2.98E-4) (**Table 6**). These convergent lines of evidence suggested CEBPB activation in aneuploid and lysosome-dysfunction cells.

**Figure 6.**
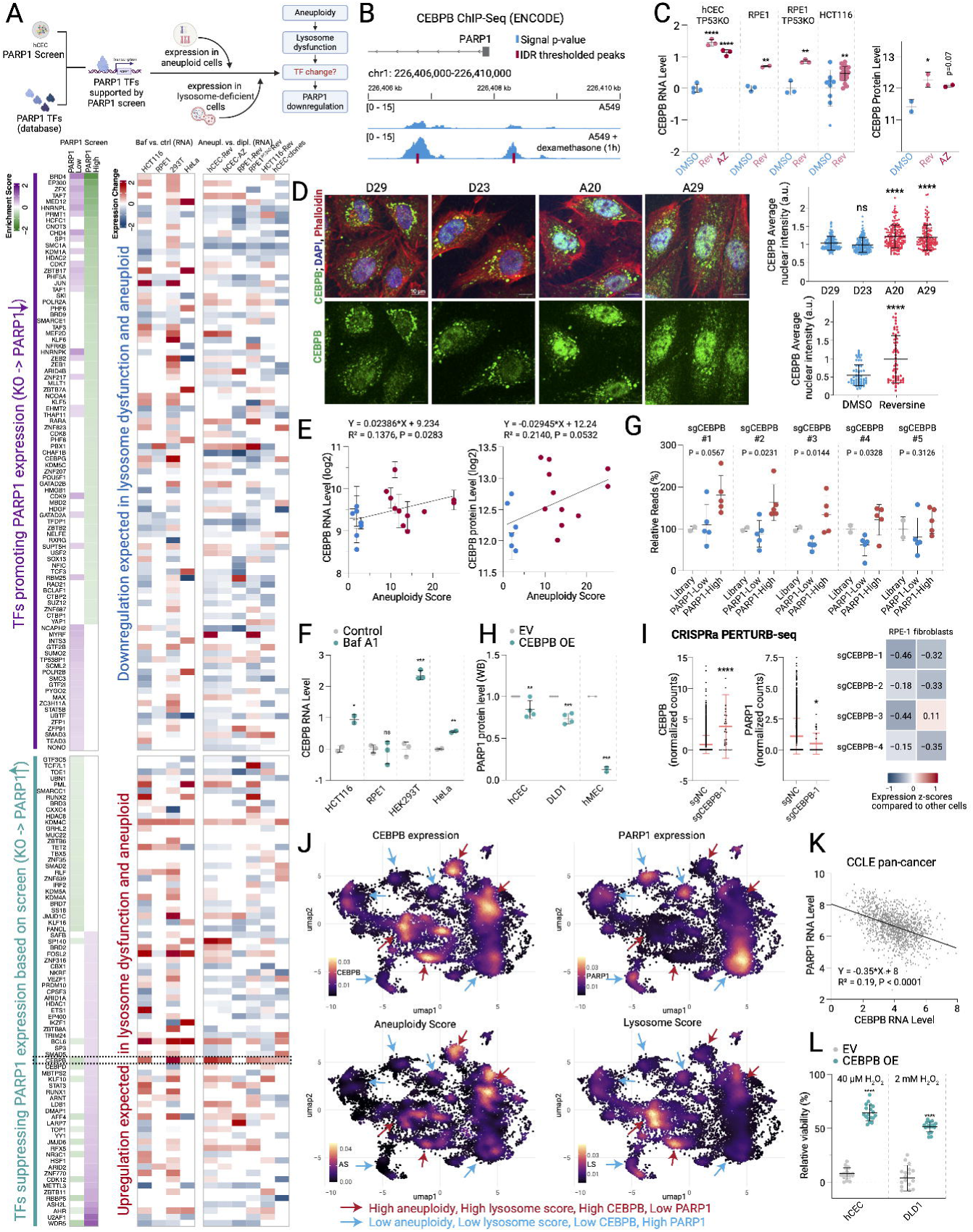
Aneuploidy-activated CEBPB transcription factor suppresses PARP1. (**A**) Top: Strategy to identify transcription factors (TFs) regulating *PARP1* expression in aneuploid cells. Bottom: Heatmap showing PARP1-promoting and PARP1-suppressing TFs and their expression changes across lysosome-deficient and aneuploid models. p<0.05. (**B**) CEBPB ChIP-Seq data (ENCODE) near the PARP1 transcription start site. (**C**) CEBPB RNA and protein levels in acute aneuploidy models. (**D**) Nuclear CEBPB in hCECs with acute and chronic aneuploidy. Left: representative images (scale bar, 10 μm). Right: quantification of nuclear CEBPB. n>50 cells/condition. (**E**) *CEBPB* RNA (left) and protein (right) correlation with aneuploidy score in aneuploid clones. (**F**) *CEBPB* RNA in Baf A1-treated cells (published dataset). (**G**) Normalized sgCEBPB read counts in sorted samples versus unsorted samples. (**H**) PARP1 protein after *CEBPB* overexpression. (**I**) CRISPR activation (CRISPRa) Perturb-Seq analysis of PARP1 regulation by CEBPB. (Left) CEBPB and PARP1 levels in cells with CEBPB CRISPRa (sgCEBPB-1) versus control (sgNC). (Right) PARP1 expression heatmap following CEBPB activation. (**J**) scRNA-seq analysis of malignant colorectal cancer cells. UMAP plots displaying cell density for cells with top 30% of CEBPB expression (upper left), top 30% of PARP1 expression (upper right), aneuploidy score >5 (lower left), and top 30% of KEGG lysosome signature score (lower right). Cell density is weighted by the respective scores. Populations with high CEBPB expression, high aneuploidy, and high lysosomal activity (red arrows) overlap but are distinct from populations with high PARP1 expression (blue arrows). (**K**) *CEBPB* and *PARP1* RNA level correlation in pan-cancer cell lines. (**L**) Viability after H_2_O_2_ in *CEBPB*-OE versus EV cells (40 μM for *TP53*-KO hCEC, 2 mM for DLD1, 24h). One-way ANOVA with Dunnett’s test (hCEC in C, G); with Tukey’s test (hCEC clones in D). Unpaired t-test (RPE1, RPE1-*TP53*KO and HCT116 in C, acute aneuploidy in D, F, H, I, L).

Given the established role of lysosomal stress in aneuploid cells^29^, we sought to further interrogate the relationship between lysosomal dysfunction and *CEBPB* activation. TFEB and TFE3 are key transcriptional regulators of lysosomal biogenesis^64^. To test whether further disrupting autophagy by inhibiting TFEB and TFE3 would exacerbate CEBPB activation, we knocked down *TFEB* or *TFE3* in diploid and aneuploid hCEC clones. Knockdown of either *TFEB* or *TFE3* significantly increased nuclear CEBPB levels specifically in aneuploid cells (A20, A21, A29) but had no significant effect in near-diploid cells (D23, D29) (**Fig. S6B**). This aneuploidy-specific response suggests that further compromising autophagy in already-saturated aneuploid cells exacerbates CEBPB activation, whereas diploid cells with intact lysosomal function are more resilient to TFEB or TFE3 disruption. Consistent with the role of CEBPB in mediating oxidative stress resistance, *TFEB* or *TFE3* knockdown significantly increased H₂O₂ resistance specifically in aneuploid clones (A20, A21, A29), with no effect observed in near-diploid cells (D23, D29) (**Fig. S6C**). These findings establish lysosomal dysfunction as an upstream trigger of CEBPB activation.

Our PARP1 CRISPR screen revealed enrichment of *CEBPB*-targeting sgRNAs in the PARP1-high population and depletion in the PARP1-low population (**Fig. 6G**), suggesting CEBPB’s suppressive role in PARP1 expression. This was further corroborated by decreased PARP1 protein upon CEBPB overexpression in various cells (**Fig. 6H**), and reduced PARP1 RNA in TCGA COREAD tumors with *CEBPB* amplification (excluding patients with PARP1 SCNA, p < 0.01, **Fig. S6D**). In a recent CRISPR activation (CRISPRa) Perturb-seq screen to evaluate the impact of transcription factor activation^65^, cells with *CEBPB* activation via CEBPB-targeting sgRNAs exhibited higher CEBPB levels and lower PARP1 levels compared to control cells (**Fig. 6I**). To directly assess whether CEBPB binds to the *PARP1* promoter, we performed chromatin immunoprecipitation followed by quantitative PCR (ChIP-qPCR). CEBPB showed significant enrichment at the *PARP1* promoter in cells treated with dexamethasone (*CEBPB* activator) and reversine compared to DMSO control (**Fig. S6E**). Collectively, these results indicate that CEBPB activation suppresses PARP1 expression.

Next, we confirmed the inverse relationship between CEBPB, PARP1, aneuploidy, and lysosomal status in human primary colorectal tumors using scRNAseq^66,67^. Cells with high CEBPB expression exhibited high aneuploidy scores, high lysosome scores, and low PARP1 expression, and vice versa (**Fig. 6J, Fig. S6F**), revealing that aneuploid cancer cells are associated with lysosomal alterations, CEBPB activation and low PARP1 expression. Consistently, we observed strong inverse correlations between CEBPB and PARP1 RNA levels across pan-cancer cell lines (**Fig. 6K**) and COREAD tumors (**Fig. S6G**). These associations across our models and human datasets, combined with the functional validation of *CEBPB* knockout or overexpression on PARP1 expression (**Fig. 6G-6I**), support CEBPB’s role as a suppressive TF of *PARP1*. In addition, CEBPB overexpression also conferred resistance to oxidative stress-induced cell death (**Fig. 6L**), mirroring phenotypes observed in aneuploid, PARP1-downregulated, and lysosome-dysfunction cells. Overall, our data establish CEBPB as a critical TF activated in aneuploid and lysosome-deficient cells, suppressing PARP1 expression and contributing to oxidative stress resistance.

## Discussion

In this study, we discovered that aneuploid cells resist ROS-induced cell death more than diploid cells (**Fig. 1, 2**). Mechanistically, *PARP1* is suppressed across 15 aneuploidy models (**Fig. 2**), and this suppression confers resistance to parthanatos, i.e., PARP1-mediated cell death, which can be triggered by ROS, DNA alkylation, or UV radiation (**Fig. 3**). Concurrently, this suppression compromises PARP1-mediated DNA damage repair in aneuploid cells (**Fig. 3**), potentially contributing to cancer progression. Accordingly, we found that PARP1 levels inversely correlate with metastatic potential in cancer models and human patients (**Fig. 4**). Through comprehensive screening, we identified lysosomal stress and subsequent CEBPB activation as potential mediators of PARP1 suppression in aneuploid cells (**Fig. 5-6**, **Fig. S6A-H**).

### Relationship between aneuploidy and ROS across different species

Earlier studies in *Saccharomyces Cerevisiae* revealed that aneuploid yeasts show increased ROS and hypersensitivity to H_2_O_2_^68^, contrasting with our findings in mammalian cells. In our model, aneuploid cell resistance to oxidative stress is not mediated by increased ROS-buffering capacity but by reduced sensitivity to ROS-induced cell death through PARP1 suppression (**Fig. 2-3**). Notably, *Saccharomyces Cerevisiae* lacks PARPs and PARP-like proteins^69^, which may explain the difference. Previous studies using *Drosophila* and mouse models with persistent chromosome instability (CIN) reported increased intracellular ROS^70,71^. Our chromosomally stable aneuploid cells showed no significant increase in ROS levels, oxidative damage, or NRF2 activation (**Fig. 2 and S2**). The discrepancy may reflect species-specific effects or differences between stable aneuploidy and ongoing CIN.

### PARP1 suppression in aneuploid cells is associated with CEBPB

PARP1 suppression in aneuploid cells does not result from cell cycle dysregulation. PARP1 downregulation occurs in both *TP53*WT and *TP53*KO aneuploid cells (**Fig. 2**), despite p53-dependent G1 arrest^72,73^. Additionally, aneuploid cells generated by different methods show PARP1 downregulation (**Fig. 2I, Fig. S2I-K**), while maintaining cell cycle profiles^74–76^ and proliferation rates similar to their diploid counterparts (**Fig. 1I**, **Fig. 4A**)

Our CRISPR screen identified PARP1 expression regulators, providing a valuable resource for PARP1 research. Integration with RNAseq data suggested that alterations in the autophagy-lysosome pathway (**Fig. S5E**), particularly lysosomal V-ATPase genes (**Fig. S5F**), may contribute to PARP1 downregulation in aneuploid cells. Lysosomal stress is a common feature in aneuploid cells (**Fig. 5F-H**)^29,30^. We demonstrated that lysosome inhibition induces PARP1 downregulation (**Fig. 5I**), conferring cell resistance to ROS-induced cell death in a PARP1-dependent manner (**Fig. 5J-K**). These results suggest lysosomal stress is a potential mediator of PARP1 downregulation in aneuploid cells.

Furthermore, we found that both aneuploidization and lysosome dysfunction lead to increased nuclear CEBPB levels (**Fig. 6**). Genetic experiments on CEBPB support its critical role as a transcriptional factor mediating PARP1 expression and ROS resistance in aneuploid cells. Existing evidence supports a potential connection between aneuploidy and CEBPB, possibly via proteotoxicity. Studies have demonstrated that autophagy activation or deficiency activates CEBPB^77–80^. Aneuploidy is known to induce proteotoxicity, which activates autophagy through transcription factor EB (TFEB)^29^. Excessive proteotoxic stress impairs autophagic flux and lysosome function. Consistently, we found that further compromising autophagy through *TFEB* or *TFE3* knockdown exacerbated CEBPB activation and enhanced ROS resistance in aneuploid cells, while having no effect on diploid cells with intact lysosomal function. Therefore, we propose that lysosome dysfunction in aneuploid cells may activate CEBPB through proteotoxicity, leading to PARP1 repression and parthanatos resistance.

### PARP1 suppression and implications for cell death, DNA repair, metastasis and cancer treatment

Parthanatos, i.e. PARP1-mediated cell death, occurs after toxic insults that cause excessive oxidative DNA damage^23,81^ but remains poorly understood in cancer^28^. PARP1 inhibition or downregulation has shown cytoprotective effects in various non-cancerous conditions^41,82–86^. Our results demonstrate that modulating PARP1 levels in diploid or aneuploid cells directly affects their resistance to parthanatos induced by cancer-related DNA damaging insults, such as ROS, DNA alkylation, and UV radiation (**Fig. 3**). These findings may explain previous observations of aneuploidy-induced resistance to certain chemotherapy and radiation therapy^10,11,87–89^.

PARP1 downregulation also impairs repair of oxidative and alkylating DNA damage in aneuploid cells (**Fig. 3M**, **Fig. S3J-K**), consistent with earlier research showing PARP1 depletion reduces SSB repair rates^90^. Aneuploid cells did not exhibit impaired repair of DNA damage induced by certain agents that may induce multiple DNA damage repair mechanisms with less dependency on PARP1. The apparently paradoxical survival advantage despite impaired DNA repair may be explained by PARP1’s dual role: excessive PARP1 activation depletes NAD+ and ATP, which can be more lethal than the DNA damage itself^26^, making PARP1 suppression pro-survival under extensive DNA damage. Taken together, we propose that aneuploidy may create a ‘perfect storm’ where resistance to cell death is coupled with decreased DNA repair under oxidative stress via PARP1 downregulation, potentially promoting tumorigenesis and cancer evolution.

PARP1 has been an important therapeutic target for HR-deficient tumors^91^. Our findings may have implications for the clinical use of PARPi. PARP1 downregulation has been associated with PARP1i resistance^92–94^, suggesting that aneuploid cancer cells may inherently resist PARP1i due to PARP1 downregulation, and analysis of CCLE cancer cell lines confirms that aneuploid cells show reduced PARPi sensitivity (**Fig. S6I**). Additionally, PARPi might counteract the cytotoxic effects of DNA damage-inducing agents if their efficacy relies on PARP1 hyperactivation and induction of parthanatos^28^. Altogether, our studies, by shedding light on a previously unappreciated effect of aneuploidy—promoting ROS resistance through PARP1 suppression—pave the way for future studies to better characterize the implications of these findings for cancer therapy and resistance.

### Limitations of the Study

Several limitations should be noted. First, as intratumoral H₂O₂ concentrations are not precisely defined, our *in vitro* models may not fully recapitulate the magnitude or duration of oxidative stress during tumor progression or treatment. Second, although we demonstrate that modest PARP1 reductions alter cellular phenotypes, systematic dose-response studies are needed to precisely define cancer cell sensitivity to graded alterations in PARP1 dosage. Third, while our *in vivo* experiments link PARP1 levels to metastatic potential, it remains unclear whether this effect operates through resistance to ROS-induced parthanatos or other mechanisms. Finally, while our data establish CEBPB as a transcriptional regulator of PARP1, the precise mechanism by which aneuploidy and lysosomal dysfunction activate CEBPB specifically in aneuploid cells warrants further investigation.

## STAR Methods

### Summary

Aneuploidy models were generated by transient MPS1 inhibitor treatment (acute model), single-cell cloning following MPS1 inhibitor treatment (chronic model), CRISPR-mediated chromosome missegregation, or microcell-mediated chromosome transfer (MMCT). Aneuploidy was confirmed by flow cytometry and whole-genome sequencing, and aneuploidy scores were calculated from arm-level somatic copy number alterations. Cell viability was assessed by fluorescence microscopy or flow cytometry following oxidative stress or other treatments. Global RNA and protein expression were evaluated by RNA-sequencing and mass spectrometry, respectively, with gene-specific detection by RT-qPCR and western blotting. Genome-wide CRISPR knockout screening was performed with FACS-based sorting by PARP1 levels, followed by sgRNA enrichment analysis using MAGeCK. PARP1, TIMELESS, CEBPB, and other genes were manipulated using lentiviral shRNA knockdown, Cas9-sgRNA knockout, or cDNA overexpression. DNA damage was assessed by alkaline comet assay and γH2AX staining. Metastasis was evaluated in NSG or B6 Albino mice by bioluminescence imaging following cancer cell injection. Integrative analysis of transcriptome and proteome data generated in this study and from published bulk and single-cell RNA-sequencing datasets was used to evaluate gene expression and pathway activity. In all figures, individual data points represent replicates from representative experiments (at least two independent experiments were performed to validate results). For dot plots, unless otherwise specified, summary statistics show mean ± standard deviation. Statistical analyses used unpaired t-tests, ANOVA with post-hoc tests, or non-parametric tests as appropriate, with p < 0.05 considered significant. Detailed methods are provided in Supplemental Information.

### Aneuploidy models

#### MPS1i-induced acute aneuploidy

To study the short-term effects of aneuploidy, irrespective of specific chromosome gains or losses, we utilized a model of **acute aneuploidy** in which near-diploid cells were treated with MPS1i (reversine or AZ3146) to induce chromosome missegregation. *TP53*WT or *TP53*KO hCECs, RPE1, DLD1, and mouse KP cancer cells were seeded at 30% confluency and treated with either 0.5 μM reversine or 5 μM AZ3146 for 24-30 hours. hMECs were treated under similar conditions but with an extended 48-hour exposure to the inhibitors. Subsequently, the cells were trypsinized and cultured in the regular medium for 3 days (the release phase). Over-confluence was avoided both during MPS1i treatment and the release phase. The control group was treated with an equivalent amount of DMSO (0.1% v/v) for an identical duration and was then released for the same time period. Both the diploid control (treated with DMSO) and the aneuploid cells (treated with reversine or AZ3146) were plated on new culture dishes or plates to achieve comparable confluency on the second day for downstream comparisons. To assess the potential side effects of MPS1i, confluent hCECs with temporarily suppressed proliferation were treated with 0.1% DMSO or MPS1i (0.5 μM reversine or 5 μM AZ3146) for 24-30 hours, followed by a 3-day release and replating for downstream experiments. Over-confluence was only avoided during the release phase. The induction of aneuploidy was confirmed through flow cytometric analysis of DNA content and scRNAseq-based CNA quantification (see description below).

#### Isogenic clones containing different aneuploidy levels

To study the longer-term effects during adaptive stages of aneuploidy, irrespective of specific chromosome gains or losses, a model of **chronic aneuploidy** was developed. Near-diploid *TP53*KO hCECs were exposed to 0.2 μM reversine for 24 hours. Then they were plated at a low density and allowed to grow until colonies formed. Single-cell-derived clones were isolated using glass cylinders and transferred to separate wells in multiwell plates for further expansion. To identify chromosomal gains and losses, the DNA of a cohort of 56 isogenic hCEC clones was extracted and subjected to shallow whole-genome sequencing (see description below). The clones were categorized as near-diploid, exhibiting 0-2 gains or losses of chromosomes or chromosome arms (designated as D followed by numbers), or as aneuploid, with ≥ 5 gains or losses of chromosomes or chromosome arms (designated as A followed by numbers). The clones maintained stable karyotypes over one month of *in vitro* culture.

#### Previously published Aneuploidy models generated by MMCT

Microcell-mediated chromosome transfer (MMCT) is a common method for generating aneuploid clones with one or two extra chromosomes^95^. Aneuploid clones of HCT116 with an extra chromosome 5 and RPE1 with an extra chromosome 7, along with their diploid control clones^32^, generously provided by Dr. Zuzana Storchová and Dr. Jason M. Sheltzer, served as an independent aneuploidy model to test our hypotheses.

### Aneuploidy score

Aneuploidy was quantified by estimating the total number of arm-level gains and losses for each hCEC clone, cancer cell line, or tumor sample. For hCEC clones, the arm-level copy number alterations were calculated based on the copy number package^96^. We considered a log2-transformed copy number ratio > 0.3 as a gain and < (–0.3) as a loss. The aneuploidy scores of CCLE cancer cell lines were downloaded from Depmap portal^97^. For Fig. 4B and 4C, cell lines were stratified into three groups based on aneuploid scores (low, medium, and high aneuploidy) with the same number of cells in each group (pan-cancer: 314 cell lines per group; COREAD: 18 cell lines per group). For TCGA COREAD samples, the arm and gene level somatic copy number alteration data were obtained from the TCGA dataset. The log2-transformed copy number ratios for each chromosomal region were adjusted by tumor purity and ploidy using ABSOLUTE method ^98^. Sample-related purity and ploidy were downloaded from TCGA Firehose Legacy. The detailed method for copy number adjustment based on purity and ploidy was described before^99^. After the adjustment, we considered a log2-transformed copy number ratio > 0.3 as a gain and < (–0.3) as a loss.

## Acknowledgments

We thank all the Davoli lab members for helpful discussions. We thank Dr. Thomas M. Norman for providing the CRISPRa Perturb-Seq data. We are grateful to Dr. Jason Sheltzer (Yale University School of Medicine) and Dr. Zuzana Storchová (TU Kaiserslautern) for the aneuploid clones generated using MMCT and their near-diploid controls. We thank Alcida Karz (NYU Grossman School of Medicine) for providing the control shRNA. We thank NYU Grossman School of Medicine’s Genome Technology Center and Proteomics Laboratory for help with experiments. This research was supported by grants from the Cancer Research UK Grand Challenge, the Mark Foundation for Cancer Research (C5470/A27144), NIH R37 R37CA248631, R01 R01HG012590, R01 R01DK135089, the MRA Young Investigator Award, the Breast Cancer Alliance Young Investigator Award and a NFCR (National Foundation for Cancer Research) and by the Cancer Center Support Grant P30CA016087 at the Laura and Isaac Perlmutter Cancer Center to NYU Langone’s Genome Technology Center (RRID:SCR_017929) and Proteomics Laboratory (RRID:SCR_017926). Biorender was used for some panels in Figures 1, S1, 5, 6 and S6.

## List of Supplementary Materials

Supplementary Materials:

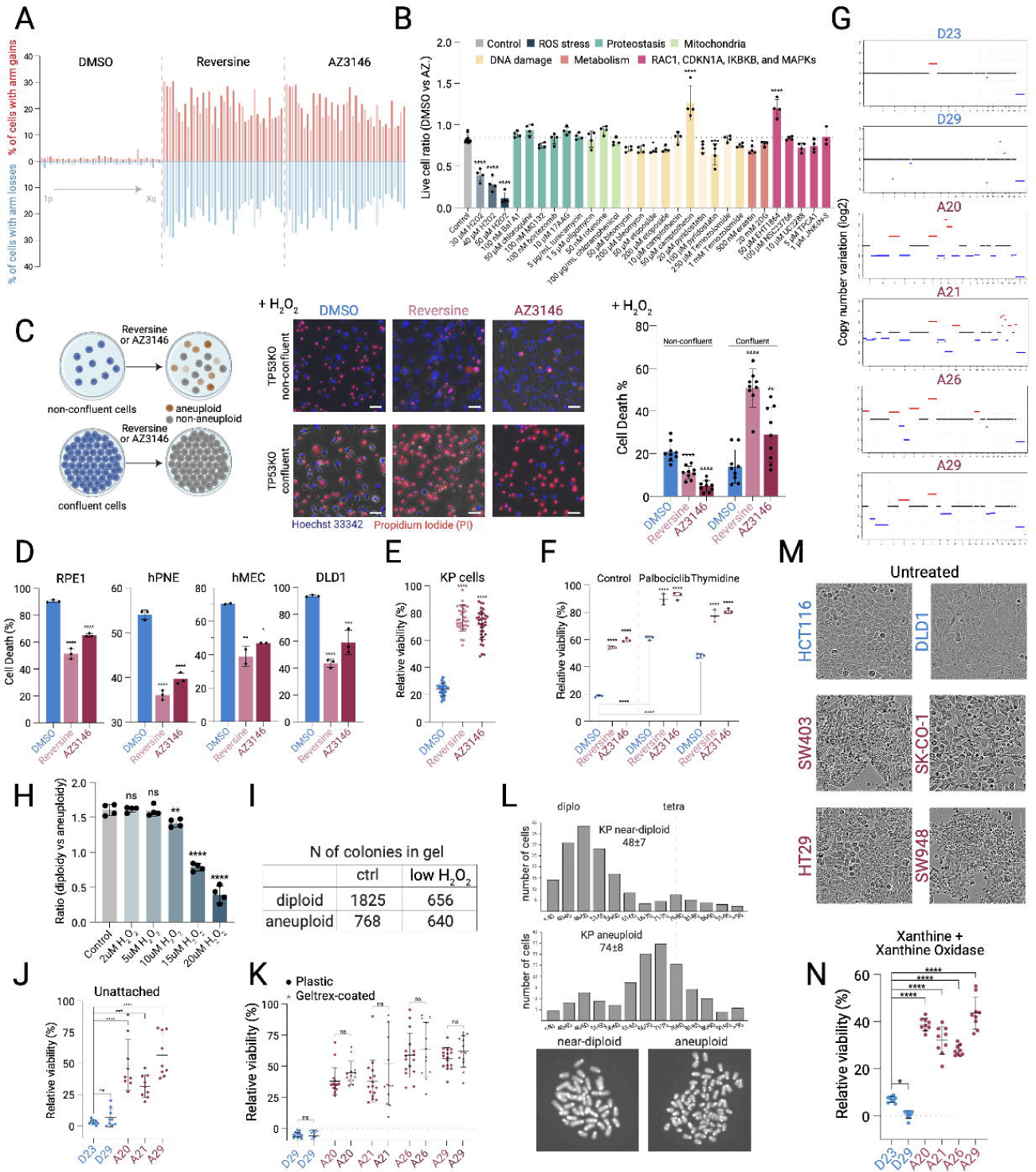

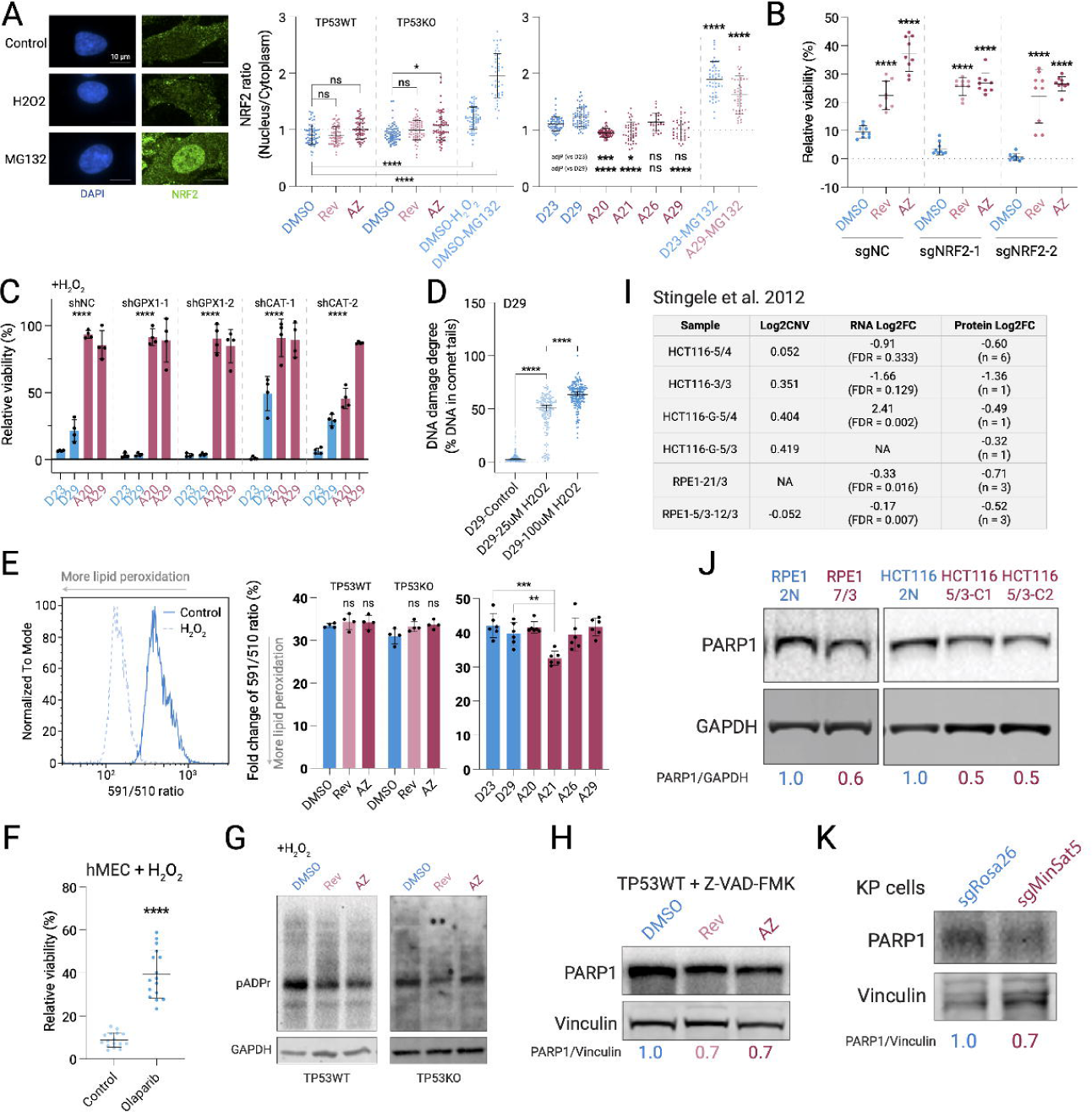

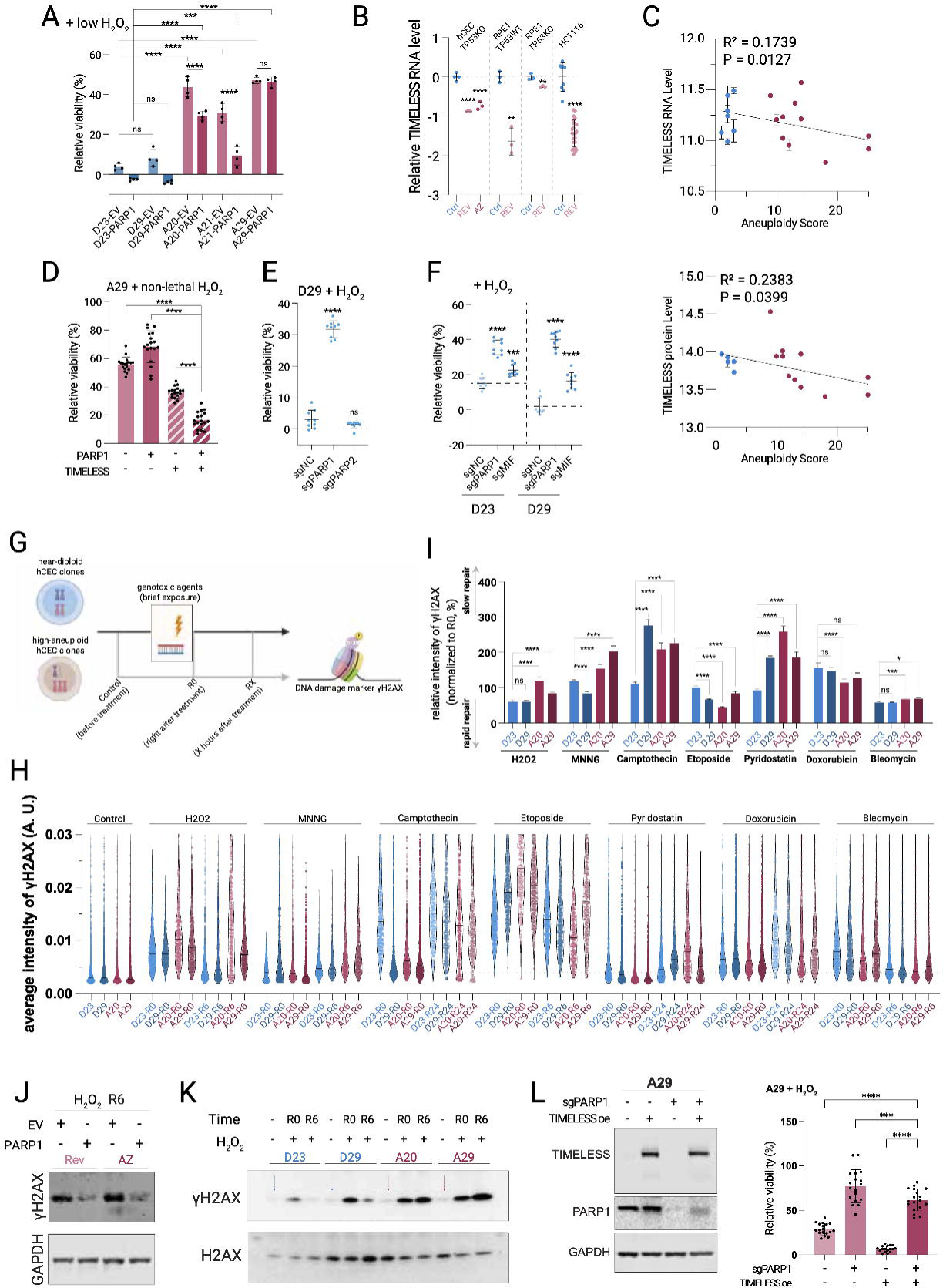

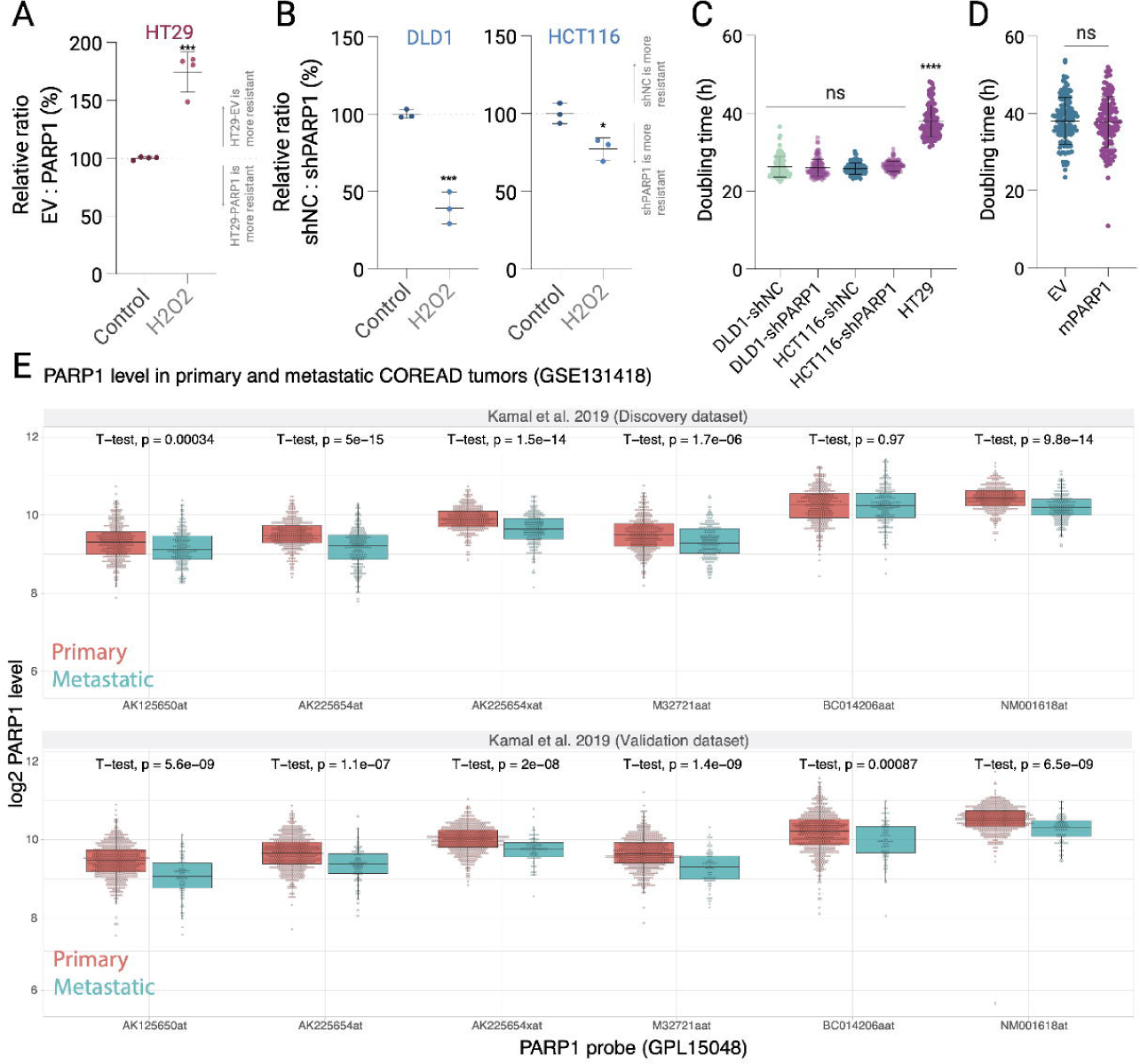

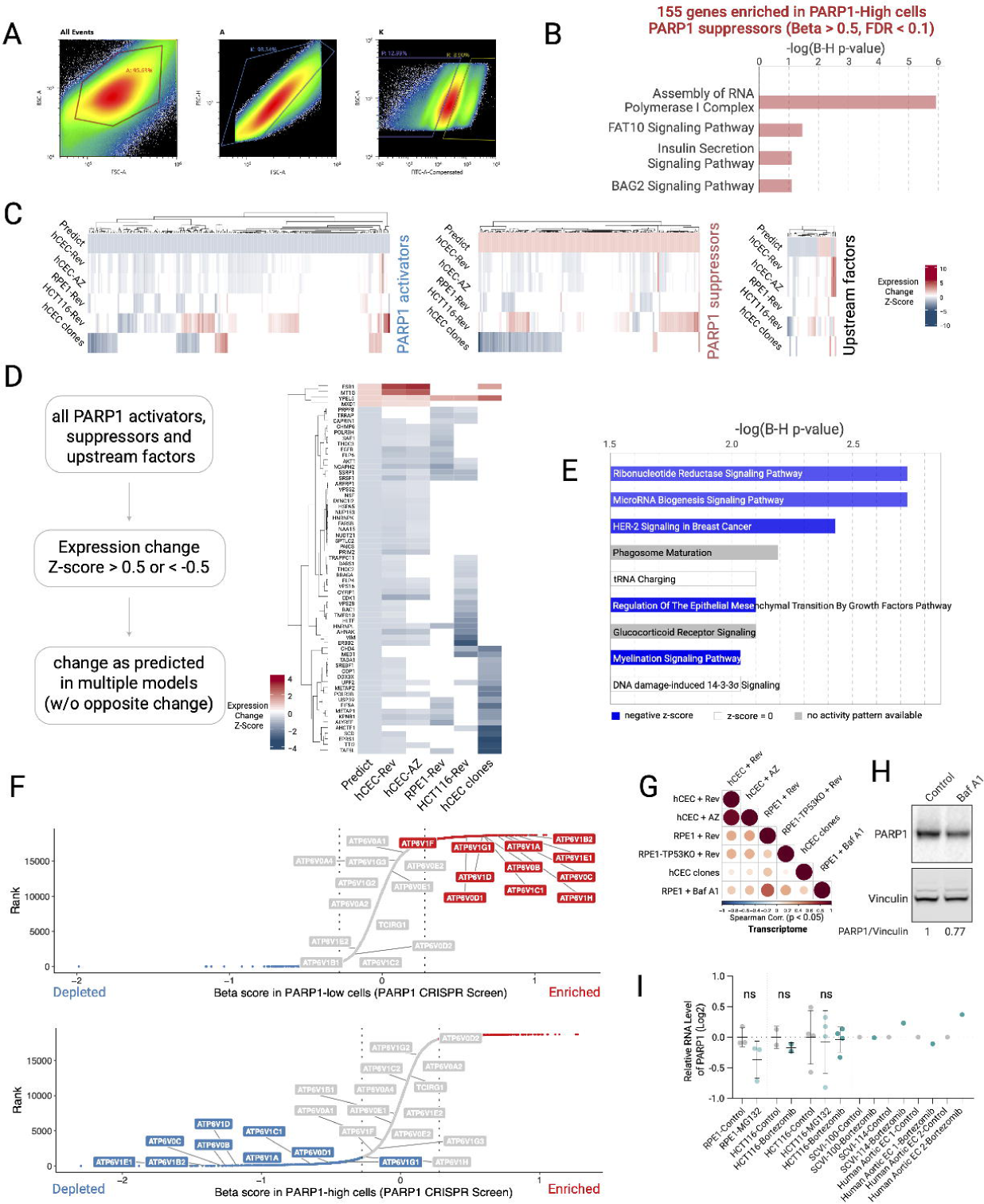

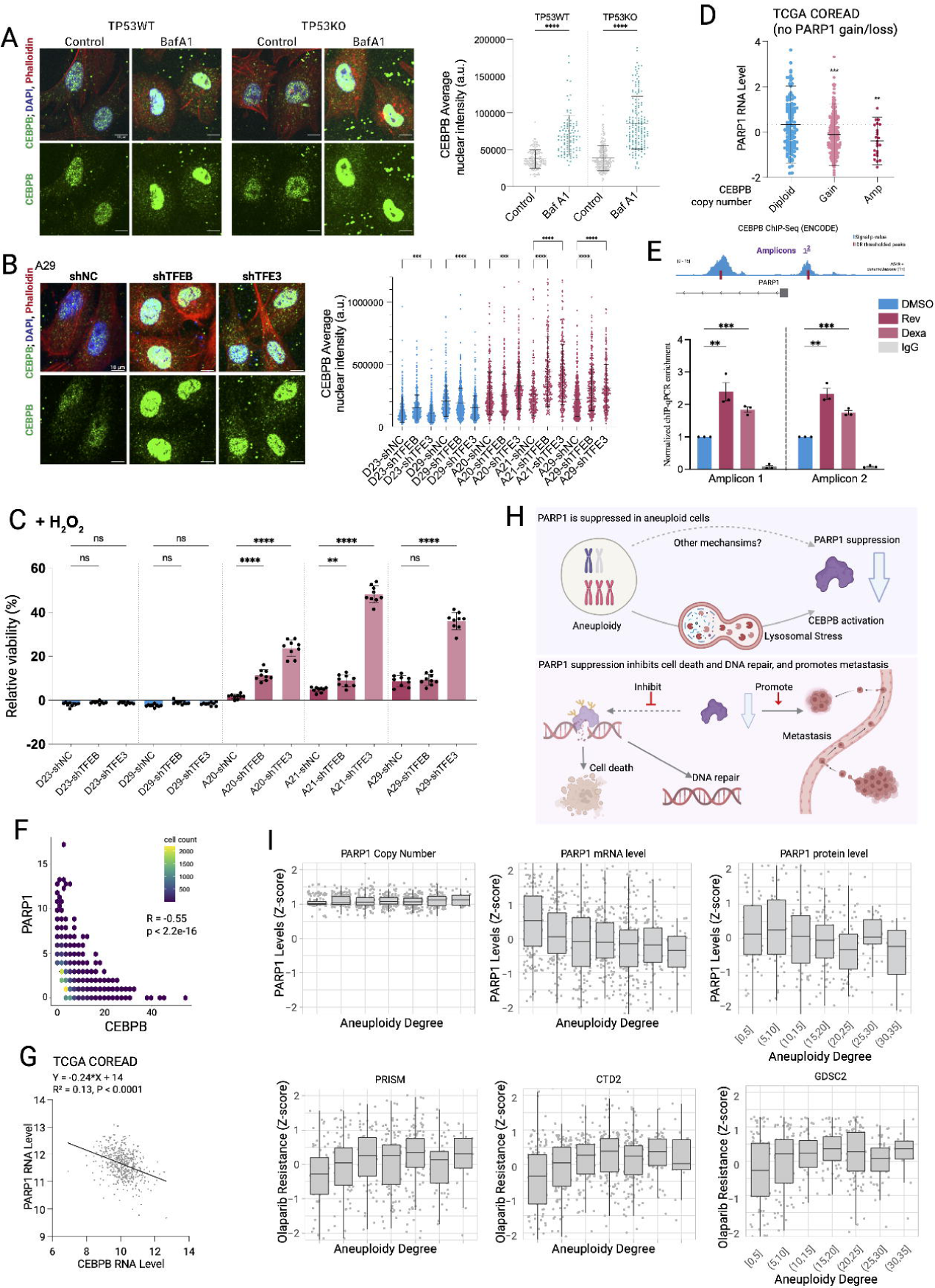

Tables S1 and S7

