## Supplemental Information for "PARP1 Suppression Drives ROS Resistance in Aneuploid Cancer Cells"

### Supplementary Information

### Detailed Methods

#### Cell Culture

hTERT-immortalized hCECs[100] were cultured in a 4:1 mixture of DMEM/Medium 199, supplemented with 2% fetal bovine serum (FBS), 5 ng/mL epidermal growth factor (EGF), 1 μg/mL hydrocortisone, 10 μg/mL insulin, 2 μg/mL transferrin, and 5 nM sodium selenite. hTERT-immortalized RPE1 (retinal pigment epithelial cells) cells were cultured in DMEM/F-12 supplemented with 10% FBS. hTERT-immortalized hPNE (human pancreatic nestin-expressing cells) were cultured in RPMI 1640 supplemented with 10% FBS. hTERT-immortalized hMEC (human mammary epithelial cells) were cultured in Mammary Epithelial Cell Culture Kit (Lonza, CC-2551B). Colon cancer cells (HCT116, DLD1, HT29, SK-CO-1, SW403, SW948), aneuploid clones of HCT116 and RPE1 cells generated through MMCT (HCT116-5/3-C1, HCT116-5/3-C2, RPE1-7/3), and their near-diploid clones (HCT116-2N, RPE1-2N), were cultured in DMEM supplemented with 10% FBS. HEK-293T were cultured in DMEM supplemented with 10% bovine calf serum (BCS). All media for the above human cells were supplemented with 100 U/mL penicillin-streptomycin and 2 mM L-glutamine. Mouse KP cancer cells were isolated from tumor nodules of the lung of KP ($Kras^{G12D}Trp53^{-/-}$) genetically engineered mouse models (GEMMs) from C57BL/6 as described previously [33,34], and cultured in RPMI 1640 supplemented with 10% FBS, 1X antibiotic-antimycotic, and 1X GlutaMax. All cells were cultured in a humidified environment at 37°C and 5% $CO_2$ and were tested free of mycoplasma contamination.

#### Plasmid construction

*TP53* was knocked out in hCECs by transfection with a Cas9-containing plasmid (Addgene #42230) and pLentiGuide-Puro expressing the following sgRNA: 5'gcatgggcggcatgaaccgg3'. Clones were derived and the effect on TP53 level was confirmed using WB.

For introduction of *PARP1*, GFP-NLS into cells, human PARP1 cDNA was PCR amplified from the pBABE-puro-Halo-PARP1[101] plasmid using the forward primer (5'caccatggcggagtcttcggataagct3') and the reverse primer (5'ttaccacagggaggtcttaaaattg3'). GFP-NLS cDNA was PCR amplified from a template plasmid using the forward primer (5'ttacaccttgcgtttttcttgggcttgtacagctcgtccatg3') and the reverse primer (5'ttaccacagggaggtcttaaaattg3'). After gel purification using the Nucleospin Gel and PCR Clean-UP kit (Macherey-Nagel, 740609), the DNA fragment was inserted into the pENTR vector using the D-TOPO cloning kit (Thermo Scientific, K240020), following the manufacturer's instructions. Then the PARP1 or GFP-NLS sequence was integrated into the pHAGE-CMV-blast or pHAGE-TRE-hygro donor vectors via the LR reaction using the Gateway cloning kit (Thermo Scientific, 11791020), following the manufacturer's protocol. For the control plasmid, the same vector was used with a short random sequence (5'GACATAGTACAG3') replacing the PARP1 or GFP-NLS sequence.

For introduction of human *TIMELESS* or mouse *PARP1* into cells, human TIMELESS cDNA was PCR amplified from a DNA template (a gift from Dr. Jef Boeke) using the forward primer (5'tacaaaaaagcaggctccgcggccgccccccttcaccatggacttgcacatgatgaa3') and the reverse primer (5'gtacaagaaagctgggtcggcgcgcccacccttctagtcatcctcatcatcct3'). Mouse PARP1 cDNA was PCR amplified from ZS-pEGFP-n-mPARP1 (a gift from Dr. Zha Shan Lab) using the forward primer (5'tacaaaaaagcaggctccgcggccgccccccttcaccatggcggaggcctc3') and the reverse primer (5'aagaaagctgggtcggcgcgcccacccttctaccacagggatgtcttaaaattgaacttg3'). The gel-purified DNA fragment was assembled with AscI/NotI-linearized pENTR vector using Gibson Assembly. The mouse PARP1 sequence was then recombined into the pHAGE-CMV-blast destination vector and the *TIMELESS* sequence into pHAGE-TRE-hygro destination vector via LR Clonase reaction.

For *PARP1*, *GPX1*, *CAT* knockdown, two shRNAs targeting each gene (shPARP1-1: cttcgttagaatgtctgcctt, shPARP1-2: gcagcttcataaccgaagatt, shGPX1-1: gcaaggtactacttatcgaga, shGPX1-2: gcatcaggagaacgccaagaa, shCAT-1: gccacatgaatggatatggat, shCAT-2: cggagattcaacactgccaat) in MISSION® TRC1 lentiviral vector were purchased from Sigma-Aldrich (TRCN0000007928 and TRCN0000007929). The MISSION® pLKO.1-puro non-mammalian shRNA plasmid targeting no known mammalian genes was used as the control (Sigma-Aldrich, SHC002).

For *PARP1*, *PARP2*, and *MIF* knockout, sgRNA sequences (sgPARP1: 5'-TAACGATGTCCACCAGGCCA-3', sgPARP2: 5'-CGGCGGCGACGGAGCACCGG-3', sgMIF: 5'-GAGGAACCCGTCCGGCACGG-3') were cloned into the LentiCRISPR-V2-FE-puro vector (Addgene #186746). The sgRNA sequences with flanking homology regions were synthesized as oligonucleotides (IDT) and inserted into BsmBI-linearized LentiCRISPR-V2-FE-puro using Gibson assembly. For NRF2 knockout, LentiCRISPR-V2-NRF2 were obtained from Addgene (186838, 186839).

For *TFEB* and *TFE3* knockdown, shRNAs targeting each gene (shTFEB: gaacaagtttgctgcccacat, shTFE3: attgttgctgacatagaatta) in MISSION® TRC1 lentiviral vector were purchased from Sigma-Aldrich (TRCN0000013111 and TRCN0000232151). The MISSION® pLKO.1-puro non-mammalian shRNA plasmid targeting no known mammalian genes was used as the control (Sigma-Aldrich, SHC002).

For the generation of aneuploidy in mouse cells, KP cells were lentivirally transduced with pINDUCER20-AuroraB-dCas9 and a control sgRNA or a gRNA targeting mouse centromeres. To clone AuroraB-dCas9 fusion protein, dCas9 without ATG and without stop codon (for N-terminal and C-terminal tagging respectively) were cloned into the pENTR vector using the D-TOPO cloning kit (Thermo Scientific, K240020). AuroraB was PCR amplified from a template vector from the ultimate ORF collection[102] using the forward primer (5'tccgcggccccttcacccatggcccagaaggagaactcc3') and the reverse primer (5'gctgtacttcttgtcctcgagcgaccctccgccggaaccgccggcgacagattgaa3'), and fused to dCas9 into pENTR-dCas9-noATG including the GGSGGGS linker. The fusion protein was then cloned into a tetracycline response element (TRE)-included donor vector, pINDUCER20 (or pIND20, neomycin resistance, doxycycline inducible promoter)[103]. The control sgRNA is 5'aacggctccaccacgctcgg3' and the centromere targeting sgRNA is 5'atctaatatgttctacagtg3'; sgRNAs were cloned into pLRCherry V2.1 as reported previously[104]. In brief, to be suitable for cloning into BbsI-digested vectors, sense oligos were designed with a CACCG 5' overhang and antisense oligos were designed with an AAAC 5' overhang and an 3' end C. The sense and anti- sense oligos were annealed, phosphorylated, and ligated into BbsI-digested pLentiGuide-Puro-FE.

**Lentivirus production and infection**

For transduction of cells, lentiviral particles were generated as follows: 15 million 293T cells per 10-cm dish were plated 24 hours before transfection. The cells were transfected with a mixture of gene transfer plasmid (6 µg) and packaging plasmids including 10 µg psPAX2 (addgene, 12260) and 5 µg pMD2.G (VSV-G; addgene, 12259), using Lipofectamine 3000 (ThermoFisher, L3000075). The medium was changed 6 hours later and virus was collected every 24 hours after transfection (three times in total) by filtering the medium through a 0.45-µm filter. Polybrene (4 µg/mL, company, cat number) was added to the filtered media before infection. The media containing lentivirus was replaced with fresh media with appropriate antibiotic selection 24 hours after infection.

**Generation of mouse aneuploid cancer cells**

We generated aneuploid mouse KP cancer cells using our recently developed method[104] to validate the findings derived from MPS1i-induced aneuploid cells. The mouse KP cancer cells were transduced to express an sgRNA and AuroraB-dCas9 to induce chromosome missegregation. Aneuploidy was confirmed through flow cytometric analysis of DNA content and chromosome counting in metaphase spreads.

**Flow cytometry-based measurement of DNA content**

One million cells were collected and resuspended in 300 µL of cold phosphate buffered saline (PBS). Then 700 µL of ice-cold 100% ethanol was added dropwise as cell suspensions were vortexed at medium speed to ensure complete fixation. The cell suspensions were stored at 4℃ overnight. On the following day, the cells were resuspended in 500 µL of staining buffer (PBS supplemented with 2 mM EDTA, 0.1 mg/mL RNAse A, 0.01 mg/mL propidium iodide (PI), and 0.5% bovine serum albumin (BSA)), and then incubated for 1-3 hours at room temperature in the dark. Then 200 µL of cell suspensions were transferred to round-bottom 96-well plates for flow cytometric measurement of cellular PI intensity. Flow cytometry acquisition was performed using the SONY SA3800 spectral cell analyzer. PI was excited by a 561 nm laser and its fluorescence was measured through the total signal of two consecutive channels centered around 617 nm. Data analysis was performed using FlowJo with consecutive gating to exclude cell debris and doublets.

**Metaphase spreading for chromosome counting**

Cells were plated in 10 cm dishes to reach 70% confluency at the time of harvest. Before harvesting, the cells were treated with 0.1 μg/mL colcemid at 37℃ for 4 hours. Then cells were trypsinized and washed once with PBS. The cell pellet was then resuspended in 0.5 mL of PBS by tapping, followed by the addition of 10 mL 0.075 M potassium chloride (KCl), pre-warmed to 37℃. After thorough mixing by gently inverting, cells were incubated for 7 minutes at 37℃. Then 1 mL of methanol-acetic acid (3:1) was added to the cell solution, followed by thorough mixing and a 10-minute incubation at room temperature. Then cells were resuspended in 0.5 mL of 0.075 M KCl, and 9 mL of methanol-acetic acid (3:1) was added dropwise while vortexing the cell suspensions at medium speed to ensure complete fixation. The fixed cells were resuspended in 0.5 mL of fresh methanol-acetic acid (3:1) and dropped from a couple of inches above the end of cold slides tilted at a 45° angle. After overnight air drying, the slides were stained with Vectashield antifade mounting media containing DAPI (H-1200-10). The mounted slides were imaged using the Invitrogen EVOS M7000 imaging system or the Nikon spinning disk confocal microscope.

**Single-cell RNA sequencing**

Single-cell RNA sequencing (scRNA-seq) was employed to infer somatic copy number alterations (SCNA) in *TP53*KO hCECs treated with either DMSO or MPS1i, as previously described[104]. scRNA-seq libraries were constructed using the 10X Chromium Single-Cell 3' v3 Gene Expression kit in accordance with the manufacturer's instructions, which also included the manufacturer's protocol for cell surface protein (hashtag antibody) feature barcoding for samples multiplexing. The CellRanger v6.1 pipeline (10X Genomics) was used to process raw scRNA-seq data, generating gene expression matrices. To identify the sample of origin for each cell barcode, HTO count data from each 10X Chromium experiment were demultiplexed using the Seurat package (v4.0.3).

After the removal of low-quality cells and potential cell doublets, a modified version of the CopyKat (v1.0.5) pipeline was used to generate a SCNA score for each chromosome arm in each cell. Hashtagged samples from the same treatment in each 10X Chromium dataset were grouped together for analysis. In each analysis, genes expressed in fewer than 5% of the cells, HLA genes, and cell-cycle genes were excluded. The log-Freeman-Tukey transformation was employed to stabilize variance, and dlmSmooth was used to smooth outliers. The DMSO-treated diploid control sample was used to calculate a baseline expression level for each gene. This value was subtracted from the samples in the set, thereby centering the expression of the control sample around 0.

Genes expressed in fewer than 10% of cells were subsequently excluded from further analysis. Then a SCNA value was generated for each chromosome arm by calculating the mean gene expression for the genes on that arm. To account for its relatively small size, a single SCNA value for the entire chromosome 18 was calculated using genes from both the p and q arms. SCNA values for chromosomes 13, 14, 15, 21, and 22 were calculated only using genes on their respective q arms. Gains or losses of a chromosome arm relative to the control sample (diploid) were called based on a threshold calculated from the control sample for each chromosome arm using the following formula:

$$threshold = median \pm (2.5 \times MAD)$$

where the median is calculated from the SCNA values for each arm in the control sample, and the median absolute deviation (MAD) is calculated by the mad function from the stats R package. A chromosome arm was recognized as gained (or lost) if its SCNA value was above (or below) the threshold for its sample set. The heatmap of arm-level SCNA was visualized using R package ComplexHeatmap (v2.6.2).

**Shallow whole-genome sequencing (WGS)**

The genomic DNA of hCEC clones was extracted from trypsinized cells using 0.3 µg/µL Proteinase K in 10 mM Tris pH 8.0 for 1 hr at 55°C, followed by heat inactivation at 70°C for 10 minutes. Genomic DNA was then enzymatically digested using NEBNext dsDNA Fragmentase for 25 minutes at 37°C, followed by magnetic DNA bead cleanup using Sera-Mag Select Beads at a beas-to-lysate ratio of 2:1 by volume. Subsequently, DNA libraries were prepared with an average fragment size of 320 bp using the NEBNext Ultra II DNA Library Prep Kit for Illumina, following the manufacturer's instructions. In brief, after phosphorylation, dA-Tailing and adaptor ligation of fragmented DNA, unique barcodes were added to each sample and PCR amplification was performed, followed by a bead cleanup. Quantification was performed using a Qubit 2.0 fluorometer and the Qubit dsDNA HS kit. The libraries were then sequenced on an Illumina NextSeq 500 at a target depth of 4 million reads in either paired-end mode (2 X 36 cycles) or single-end mode (1 X 75 cycles). Low-pass (~0.1-0.5X) WGS reads of hCEC were aligned to reference human genome hg38 using BWA-mem (v0.7.17) and the duplicates was removed using GATK (Genome Analysis Toolkit, v4.1.7.0) to generate analysis-ready BAM files. The resulting BAM files were processed using the R package CopywriteR (v1.18.0) to determine the arm-level copy numbers. The heatmap of arm-level SCNA was visualized using R package ComplexHeatmap (v2.6.2).

**Assessing cellular sensitivity to different types of cellular perturbations**

Cell sensitivity to different cellular stress between diploid and aneuploid hCECs was compared. First, *TP53*WT diploid hCECs were labeled with nuclear-localized GFP using lentiviral transduction, followed by fluorescence-activated cell sorting (FACS). Then the unlabeled hCECs were treated with MPS1i to induce aneuploidy, while the labeled hCECs were treated with DMSO as a diploid control.

One day before the addition of the stress-inducing drugs, DMSO-treated diploid hCECs (labeled with GFP) and MPS1i-treated aneuploid hCECs (unlabeled) were plated into individual 96-well plates at a density that allowed cells to reach a comparable confluent level of 60% when cells were treated with drugs. On the second day, the culture media was replaced with fresh media containing one of the following drugs: 30-50 μM $H_2O_2$, 100 nM bafilomycin A1, 50 μM chloroquine, 100 nM MG132, 100 nM bortezomib, 10 μM 17AAG, 5 μg/mL tunicamycin, 1.5 μM oliomycin, 50 nM rotenone, 100 μg/mL chloramphenicol, 50 or 200 μM bleomycin, 50 or 200 μM etoposide, 10 or 50 μM camptothecin, 20 or 100 μM pyridostatin, 250 or 1000 μM temozolomide, 500 nM erastin, 20 mM 2DG, 50 μM EHT1864, 100 μM NSC23766, 10 μM UC2288, 5 μM TPCA1, or 1 μM JNK-IN-8. The cells were grown in the presence of the drugs for 24 hours. Following this, the cells were harvested and the labeled diploid hCECs were mixed with the unlabeled aneuploid hCECs under identical conditions. The cell mixture was stained with 2 μg/mL PI for 10 minutes in the dark, and then analyzed using flow cytometry to assess the cell number ratios between viable diploid (PI- and GFP+) and aneuploid (PI- and GFP-) hCECs under varying conditions.

**Assessing cellular sensitivity to oxidative stress**

To compare $H_2O_2$ sensitivity, cells were cultured on multiwell plates to achieve a comparable confluence at the time of $H_2O_2$ addition. On the second day, the culture medium was replaced with fresh medium containing $H_2O_2$. Cells were subjected to a gradient of $H_2O_2$ concentrations to determine the concentrations at which a portion of cells died, while avoiding high concentrations that resulted in complete cell death for all cells. Following a 24-hour treatment in a humidified environment at 37℃ with 5% $CO_2$, the relative cell viability was examined (see description below). In some cases, cells were treated with $H_2O_2$ for one hour and then released for 24 hours before the detection of the relative viability. The brief $H_2O_2$ exposure aimed to mitigate the influence of altered cell cycles and varying proliferation rates in diploid and aneuploid cells. The figure legends specify the concentration and duration of $H_2O_2$ treatment.

To assess cell sensitivity to oxidative stress induced by lower concentrations of $H_2O_2$ over an extended duration, we used two independent assays to compare near-diploid and aneuploid hCEC clones. First, three GFP-labeled near-diploid hCEC clones and three unlabeled aneuploid hCEC clones were mixed in equal numbers. These mixed cells were cultured in the presence of low concentrations of $H_2O_2$ for three days (2-20 μM), without being confluent. The ratio of live cell numbers between GFP-labeled near-diploid and unlabeled aneuploid hCECs were measured using flow cytometry. Second, 10,000 near-diploid and aneuploid hCECs, with each group consisting of three hCEC clones with different karyotypes in equal cell numbers, were mixed with 2 mL of 37℃ 0.45% low melting temperature agarose, with or without 10 μM $H_2O_2$, and added to 6-well plates pre-coated with 0.9% low melting temperature agarose. After the gel solidified, the cells were grown in a humidified environment at 37℃ with 5% $CO_2$, and fresh medium with or without 10 μM $H_2O_2$

were added to the top of the agarose daily. After ten days of growth, the colonies were stained with 150 μL of 2 mg/mL nitro blue tetrazolium and imaged with the Microtek scanner (TMA 1600-III, ScanMaker 9800XL plus). The number of colonies for near-diploid and aneuploid hCECs were quantified using Fiji software and compared.

To evaluate cell sensitivity to oxidative stress induced by alternative methods, we compared the relative viability between near-diploid and aneuploid hCEC clones after treatment with 100 μM xanthine and 20 mU/mL xanthine oxidase for 24 hours. Control cells were treated with 100 μM xanthine alone for the same duration. In the presence of the xanthine and xanthine oxidase, both superoxide and $H_2O_2$ are generated. Cells were subjected to varying concentrations of xanthine and xanthine oxidase to determine the concentrations at which a portion of cells died, while avoiding high concentrations that resulted in complete cell death for all cells.

To compare the $H_2O_2$ sensitivity of unattached cells (Fig. S1J), 100 μL of medium with or without 45 μM $H_2O_2$ was pre-added to 48-well plates. Then 35,000 hCECs in 200 μL of medium were added into the 48-well plates, followed by thorough mixing. Cell viability was examined after a 24-hour culture in a humidified environment at 37℃ and 5% $CO_2$. To compare the $H_2O_2$ sensitivity of cells attached to different surfaces (Fig. S1K), multiwell plates were coated with cold DMEM containing 1% GelTrex (LDEV-free reduced growth factor basement membrane matrix) in a humidified environment at 37℃ and 5% $CO_2$ for at least one hour. After removing the coating solution, cells were plated on the multiwell plates to achieve a comparable confluence at the time of $H_2O_2$ addition. 50 μM $H_2O_2$ were added into plates on the second day and cell viability was examined 24 hours after $H_2O_2$ treatment.

**Sensitivity to other drugs or UV treatment**

To arrest diploid cells at the G1 or S phase, *TP53* KO diploid hCECs were treated with 0.5 μM palbociclib (a CD4/6 inhibitor, Selleck Chemicals, S1116) or 2 mM thymidine (a DNA synthesis inhibitor, Sigma-Aldrich, T1895) for 24 hours. Following this treatment, the culture medium was replaced with a fresh medium containing 100 μM $H_2O_2$ in addition to 0.5 μM palbociclib or 2 mM thymidine for another 24 hours to compare the ROS resistance of the arrested cells.

To determine the specific form of cell death activated under oxidative stress, we treated the cells with inhibitors of distinct cell death pathways, including apoptosis (10 μM Z-VAD-FMK, Selleck Chemicals, S7023), necrosis (20 μM necrostatin-7, Cayman Chemical, 10528), ferroptosis (20 μM ferrostatin-1, Sigma-Aldrich, SML0583), and parthanatos (10 μM olaparib, Cell Signaling, 93852S), before and after $H_2O_2$ treatment to test if any of them was able to rescue the effects of $H_2O_2$ treatment on diploid clones. The diploid hCEC clone or hMEC cells was pretreated with the cell death inhibitors for 2 hours, followed by a one-hour exposure to 50 μM $H_2O_2$ without the inhibitors.

Then the cells were released from $H_2O_2$ and incubated with cell death inhibitors for 24 hours before the detection of cell viability.

To examine the role of NAD+ depletion during PARP1-mediated cell death, the hCEC clones were pre-cultured in medium with and without the supplementation of 180 μM NMN (Sigma-Aldrich, N3501) for 24 hours before $H_2O_2$ treatment.

To compare cell resistance to alkylating agents, near-diploid and aneuploid hCEC clones, either with or without PARP1 overexpression, were treated with 62.5 μM MNNG (Selleck Chemicals, E0157) for 24 hours before the detection of cell viability.

To compare cell resistance to UV radiation, the hCEC clones were plated on 6-well plates to reach a comparable confluent level of 60% when cells were irradiated with UV. On the second day, the cells were washed with PBS, followed by UV irradiation (50 J/m2, cells were in PBS) with the lid removed (UV Stratalinker 2400). Then PBS was replaced with fresh media and cells were cultured in a humidified environment at 37℃ and 5% $CO_2$ for 5 days. The confluent cells were split during this period. After the 5-day release, the cell number ratios of UV-treated cells to untreated cells (the relative viability) for each clone were compared.

To assess DNA damage repair in diploid and aneuploid cells, two near-diploid (D23 and D29) and two high-aneuploidy (A20 and A29) hCEC clones were transiently exposed to various genotoxic agents to induce DNA damage through distinct mechanisms, including $H_2O_2$ (50 μM, 1 hour), MNNG (125 μM, 1 hour), camptothecin (Selleck Chemicals, S1288, 10 μM, 2 hours), etoposide (Selleck Chemicals, S1225, 200 μM, 1 hour), pyridostatin (Selleck Chemicals, S7444, 20 μM, 2 hours), doxorubicin (Selleck Chemicals, S1208, 10 μM, 2 hours) and bleomycin (Selleck Chemicals, S1214, 50 μM, 1 hour).

To examine the effect of lysosome inhibitor Baf A1 on PARP1 expression and cell resistance to $H_2O_2$, hCECs (*TP53*WT and *TP53*KO) were incubated with 5 nM Baf A1 for 1 to 24 hours and then collected for WB, or treated with 200 μM $H_2O_2$ for 24 hours to compare their resistance to $H_2O_2$.

**Brightfield and widefield fluorescence imaging**

The brightfield images of cells before and after $H_2O_2$ treatment were acquired using the EVOS M5000 imaging system. To distinguish between live and dead cells using fluorescence imaging, the cells were first stained with 2.5 μM SYTO 13 (or 1 μg/mL Hoechst 33342) and 3 μg/mL PI for 15 minutes in the dark. Then they were imaged using the Incucyte S3 system (for SYTO 13 and PI) or the Cytation Cell Imaging Reader (for Hoechst 33342 and PI). In the case of the Incucyte S3 system, the signal from SYTO 13 and PI was acquired using the Green cube (441-481 nm Ex; 503-

544 nm Em) and the Red cube (567-607 nm Ex; 622-704 nm Em), respectively. For the Cytation Cell Imaging Reader, the signal from Hoechst 33342 and PI was acquired using the DAPI cube (377 nm Ex; 447 nm Em) and the Texas Red cube (586 nm Ex; 647 nm Em), respectively. Live cells were identified as being SYTO 13 (or Hoechst 33342) positive and PI negative, while dead cells were identified as being SYTO 13 (or Hoechst 33342) positive and PI positive. The brightness, contrast and size of the images were adjusted using Fiji software.

**Measurement of relative cellular viability**

To assess the relative viability of cells under drug treatment, two independent methods based on fluorescence imaging or flow cytometry were used. The relative viability (%) was calculated using the formula:

$$relative\ viability\ (\%)\ = \frac{Number\ of\ live\ cells\ under\ treatment}{Number\ of\ live\ cells\ without\ treatment} \times 100$$

To measure relative viability through fluorescence imaging, cells with and without treatment were stained with 1 μg/mL PI and 2.5 μM SYTO 13 for 15 minutes in the dark. The brightfield and fluorescence images of cells under different conditions were acquired using the Incucyte S3 system (brightfield channel, GFP channel for SYTO 13, and RFP channel for PI). Live cells were identified as being SYTO 13 positive and PI negative (GFP+ and RFP-), while dead cells were identified as being SYTO 13 positive and PI positive (GFP+ and RFP-). The counting of GFP+ and RFP+ cells for each image was performed using CellProfiler software. The number of live cells was determined by subtracting the count of RFP+ cells from the count of GFP+ cells. For mouse KP cancer cells, the cell confluence in the brightfield channel calculated by Incucyte S3 software was used as a representation of the number of live cells because the cells were too close to each other to accurately count the number of stained cells.

To measure relative viability through flow cytometry, cells with and without treatment were harvested by trypsinization. A known number of GFP+ cells were spiked into each sample as a reference, and cells were subsequently stained with 1 μg/mL PI for 10 minutes in the dark. The cell mixture was then analyzed using flow cytometry (SONY SA3800) to determine the cell number ratio of GFP- and GFP+ cells with exclusion of PI positive dead cells. The number of live cells for each condition was calculated using the following formula:

$$N\ of\ live\ cells\ = \frac{\%\ of\ GFP^{-}\ sample\ cells\ detected\ by\ flow\ cytometry}{\%\ of\ GFP^{+}\ reference\ cells\ detected\ by\ flow\ cytometry} \times N\ of\ GFP^{+} reference\ cells\ spiked\ in$$

In Fig. S1C and S1D, the percentage of dead cells was used to represent the cell sensitivity to the treatment. To achieve this, cells were stained with 1 μg/mL PI and 1 μg/mL Hoechst 33342 for 15 minutes in the dark. Then the counting of live cells (Hoechst 33342 positive and PI negative) and dead cells (Hoechst 33342 positive and PI positive) was based on flow cytometry using SONY SA3800 or fluorescence imaging using Cytation Cell Imaging Reader.

**Cell proliferation assay**

To assess cell proliferation rate, cells were plated in multiwell plates to reach a confluent level of 20% on the second day. The second day, the cells were provided with extra fresh medium, followed by the acquisition of brightfield images at regular intervals (every 4-6 hours) from multiple non-overlapping planes of view for each well using Incucyte S3 (10x objective) until the cells reached complete confluence. Cell confluence was quantified using the Incucyte basic quantification function. The confluence data was further processed using R studio to exclude confluence values below 20% or above 80%. The resulting confluence data was fitted using the GraphPad Exponential (Malthusian) growth model as follows:

$$Y = Y_0 \times e^{k \times t}$$

Doubling time, calculated as $ln(2)/k$, is the time needed for the cell population to double.

**Alkaline comet assay**

DNA damage at the single-cell level, both before and after $H_2O_2$ treatment, including single-strand and double-strand DNA breaks, and apurinic and apyrimidinic sites, was detected by the alkaline comet assay. The assay principle relies on the ability of denatured cleaved DNA fragments to migrate out of the cell under the impact of an electric potential, while undamaged supercoiled DNA remains confined within the cell membrane when a current is applied. To perform this assay, cells, both with and without one-hour exposure to 50 µM $H_2O_2$, were collected by trypsinization, washed with cold PBS, and resuspended in cold PBS at a concentration of 100,000 cells/mL. Then 40 µL of the cell suspensions were mixed with 400 µL of molten 1% low melting temperature agarose (at 37°C), and 40 µL of this mixture was immediately dropped onto 8-well hydrophobic printed slides (Electron Microscopy Sciences, 63422-11) which were pre-coated with 1% normal melting temperature agarose. The slides were left for 10 minutes at 4°C in the dark to allow the gel to solidify. Then the slides were immersed in pre-chilled lysis buffer (comprising 2.5 M NaCl, 100 mM EDTA, 10 mM Tris Base, 1% sodium lauryl sarcosinate, and 1% Triton X-100) for 45 minutes at 4°C in the dark, followed by immersion in a fresh alkaline solution, pH > 13, (consisting of 300 mM NaOH and 1 mM EDTA) for 45 minutes at room temperature in the dark. The slides were then transferred to a horizontal electrophoresis apparatus with the cold alkaline solution covering them. Electrophoresis was performed for 30 minutes at 4°C with a voltage set at about 1 V/cm x distance between electrodes (buffer volume was adjusted until the current was approximately 300 mA). Then the slides were immersed in 70% ethanol for 5 minutes. After overnight air drying, the slides were stained with SYBR Gold nucleic acid gel stain, rinsed with PBS, followed by confocal imaging (using a x4 objective and GFP channel of a Nikon spinning disk confocal microscope). The relative DNA contents in comet tails compared to the whole comets, quantified using CellProfiler software, was used to represent the degree of DNA damage.

**Detection of lipid peroxidation**

To detect degrees of lipid peroxidation with or without $H_2O_2$ treatment, C11-BODIPY 581/591 (Thermo Fisher, D3861), a lipid-soluble ratiometric fluorescent probe of lipid oxidation, was used to stain hCECs (5 μM for 45 minutes at 37℃ in the dark). The probes were integrated into the cellular membranes, mimicking the properties of natural lipids. Oxidation of the polyunsaturated butadienyl portion of the probes lead to a shift of the fluorescence emission peak from ~590 nm to ~510 nm, providing a method to measure cell antioxidant activity in lipid environments. After staining, the cells were washed with PBS and exposed to 50 μM $H_2O_2$ (or blank control) in fresh media for 30 minutes. Then the cells were collected by trypsinization and the fluorescence emission ratio of 591 nm to 510 nm emitted by the probes was measured using flow cytometry.

**Immunofluorescence**

Coverslips were coated with cold DMEM containing 1% GelTrex (LDEV-free reduced growth factor basement membrane matrix) in a humidified environment at 37℃ with 5% $CO_2$ for at least one hour. Then the cells were plated on these pre-coated coverslips in 24-well plates. On the second day, the cells were treated with drugs indicated in the results and legends. After the treatment, the cells were washed with PBS, and then fixed in 4% paraformaldehyde solution in PBS for 10 minutes at room temperature and washed with PBS for 10 minutes. Fixed cells were permeabilized with 0.5% Triton X-100 in PBS for 20 minutes at room temperature and washed with PBS for 10 minutes. The cells were then blocked with the blocking solution (1 mg/mL BSA, 3% goat serum, 0.1% Triton X-100, and 1mM EDTA at pH 8.0, in PBS) for 1 hour at room temperature and washed with PBS twice. Then the cells were incubated with primary antibodies (see below) diluted in the blocking solution at 4℃ overnight, followed by three washes with PBS. Negative controls, used to assess nonspecific background, underwent a similar process but were incubated with the blocking solution without primary antibodies. Next, the cells were incubated with secondary antibodies (see below) diluted in the blocking solution for two hours at room temperature, followed by three washes with PBS. In case of Phalloidin staining, the cells were next incubated with DyLight 554 Phalloidin (Cell Signaling Technology, 13054) diluted 1:400 in PBS for 15 minutes, followed by PBS washing. The coverslips were mounted with Vectashield antifade mounting media containing DAPI (H-1200-10). Images were acquired using the Nikon spinning disk confocal microscope at a 100x objective, ensuring overexposure was prevented during imaging. The ratio of area-normalized NRF2 fluorescence intensity in the nuclei to that in the cytoplasm, the area-normalized γH2AX fluorescence intensity in the nuclei, and the area-normalized CEBPB fluorescence intensity in the nuclei (ratio of the total intensity to the area for each nucleus) were quantified using CellProfiler software. The NRF2 primary antibody (Proteintech, 16396-1-AP) was diluted at 1:200, γH2AX primary antibody (Sigma, 05-636) was diluted at 1:1000, and CEBPB primary antibody (Abcam, ab32358) was diluted 1:100. Invitrogen goat anti-rabbit (A11034) IgG secondary antibody conjugated with Alexa Fluor 488 was used for NRF2 and CEBPB primary antibodies and anti-mouse (A28175) IgG secondary antibody conjugated with Alexa Fluor 488 was used for γH2AX primary antibody, at a dilution of 1:1000.

**Western blot**

A total of 1,000,000 cells were harvested by trypsinization, washed with cold PBS, and lysed in 100 μL of RIPA lysis buffer (Abcam, ab156034) supplemented with the protease and phosphatase inhibitor cocktail (Thermo Scientific, 78441). To detect pADPr, the cells were treated with 2 μM PDD 00017273 (a cell-permeable inhibitor of Poly(ADP-ribose) Glycohydrolase) for 1 hour before and during $H_2O_2$ treatment to suppress pADPr degradation. Then an equivalent number of cells were directly lysed in plates on ice using a specialized lysis buffer containing the inhibitors of PARP and PARG due to the rapid dynamic of PARP1 synthesis and degradation even after cell lysis [105]. This lysis buffer consisted of 50 mM Tris-HCl, pH 8.0, 100 mM NaCl, 1% Triton X-100, 5 mM MgCl2, 5 mM Bond-Breaker TCEP solution, 1x protease and phosphatase inhibitor cocktail (Thermo Scientific, 78441), 1.6 μM ADP-HPD (Sigma-Aldrich, 118415), 1 μM olaparib (Cell Signaling, 93852), and 125 U/mL benzonase (Sigma-Aldrich, E1014).

The lysates were agitated for 30 minutes at 4℃, followed by high-level sonication for 5 minutes at 4℃ (Diagenode Bioruptor Standard Sonication System, UCD-200). Then the lysates were centrifuged at 4°C and the supernatants were collected. Subsequently, the Pierce LDS buffer (Thermo Scientific, 84788) and the Bond-Breaker TCEP solution (Thermo Scientific, 77720) were added into the supernatants to reach concentrations as recommended by the manufacturers. The lysates were then boiled for 10 minutes at 80°C. After returning to room temperature, 15 uL of the lysates were loaded into wells of NuPAGE 4-12% Bis-Tris mini gels (Thermo Scientific, NP0323BOX) for electrophoresis, followed by transfer to either PVDF membranes (Bio-Rad, 1704274) using a semi-dry system (Trans-Blot Turbo transfer system, Bio-Rad, 1704150), or to nitrocellulose membranes (Bio-Rad, 1620212) using a tank transfer system (Mini Trans-Blot Cell, Bio-Rad, 1703930), following the manufacturer's protocols. The membranes were then blocked with 5% non-fat milk in Tris-buffered saline (TBS) with 0.1% Tween-20 (TBST) for 1 hour at room temperature, followed by an overnight incubation with primary antibodies diluted in 2.5% BSA in TBS (see below) at 4°C. Subsequently, the membranes were washed three times with TBST and incubated with either fluorophore- or horseradish peroxidase (HRP)-conjugated secondary antibodies diluted in 2.5% BSA in TBST for 2 hours at room temperature. Then the membranes were washed three times with TBST before signal detection. For fluorophore-conjugated secondary antibodies, the signal was directly detected using the Li-Cor Odyssey DLx imaging system. For HRP-conjugated secondary antibodies, the membranes were incubated with Pierce ECL Western Blotting Substrate (Thermo Scientific, 32106) in the dark for three minutes and then imaged using the Bio-Rad Gel Dox XR imaging system. Quantification of WB images was performed using Image J by calculating the signal derived from the indicated WB bands and subtracting the relative background. Normalization was performed relative to the indicated internal control protein. In the case of Reversine or AZ3146 treated cells, normalization to the total percentage of aneuploid cells was performed.

The following antibodies and dilutions were used: TP53 (Santa Cruz, sc-126) at a dilution of 1:100, PARP1 (Proteintech, 13371-1-AP or Cell Signaling, 9542T) at a dilution of 1:1000, pADPr (Sigma-Aldrich, MABE1031) at a dilution of 1:1000, γH2AX (Sigma-Aldrich, 05-636) at a dilution of 1:1000, H2AX (Cell Signaling Technology, 2595) at a dilution of 1:1000, TIMELESS (ab109512) at a dilution of 1:1000, GAPDH (Santa Cruz, sc-47724) at a dilution of 1:10,000, vinculin (Sigma-Aldrich, V9131) at a dilution of 1:20,000, IRDye 680RD goat anti-mouse IgG secondary antibody (926-68070) at a dilution of 1:20,000, IRDye 800CW goat anti-rabbit IgG secondary antibody (926-32211) at a dilution of 1:20,000, HRP-conjugated goat anti-mouse IgG secondary antibody (ab205719) at 1:10,000, HRP-conjugated goat anti-rabbit IgG secondary antibody (ab205718) at 1:10,000.

**RNA sequencing and data processing**

Cells were plated on 6-well plates one day before the collection. On the second day, the cells were examined to ensure their confluency level between 70% to 90%, with normal morphological characteristics. Then the cells were washed twice with PBS and promptly stored at –80°C. Total RNA was extracted using the PicoPure RNA Isolation kit (Life Technologies, Frederick, MD), which included an on-column RNase-free DNase I treatment (QIAGEN, Hilden, Germany), following the manufacturer's recommendations. To further purify the RNA for sequencing, the QIAGEN RNeasy Mini Kit (QIAGEN, 74106) was used. The concentration and integrity of the RNA were assessed using a 2100 BioAnalyzer (Agilent, Santa Clara, CA). Subsequently, sequencing libraries were constructed using the TruSeq Stranded Total RNA Library Prep Gold mRNA kit (Illumina, San Diego, CA) with an input of 250 ng RNA and a 13-cycle final amplification. The resulting libraries were quantified using High Sensitivity D1000 ScreenTape on a 2200 TapeStation (Agilent) and the Qubit 1x dsDNA HS Assay Kit (Invitrogen, Waltham, MA). The samples were equimolarity pooled, and sequencing was carried out on an Illumina NovaSeq6000 SP 100 Cycle Flow Cell v1.5 in paired-end 50 reads mode.

The total RNA-sequencing reads were mapped to the human genome hg38 using STAR (v2.7.7a)[106] with a 2-pass model. The hg38 reference sequence and RefSeq annotation were obtained from the UCSC table browser. The quantification of gene and transcript expression levels was performed using RSEM (v1.3.1)[107], which generated gene-level raw counts and fragments per kilobase of transcript per million mapped reads (FPKM) results in table format. The RNA RSEM data was filtered to include genes with a median FPKM > 1 for downstream analyses. The expression change (Log2FC) of reversine- and AZ3146-treated hCECs compared to DMSO-treated hCECs were calculated using DESeqDataSetFromMatrix and DESeq functions in the R package DESeq2 (v1.34.0)[108], which was used in Fig. 2D, 6C, and Fig. S3B, with the mean for the control being set to zero. For hCEC clones with different degrees of aneuploidy, the raw counts table was normalized using DESeqDataSetFromMatrix, estimateSizeFactors and counts functions in the R package DESeq2, followed by Log2 transformed. The log2-transformed normalized RNA levels were fitted by aneuploidy degrees of hCEC clones with a linear model as the following formula:

$$RNA\ log2FC \sim \beta_0 + \beta_1\ \times\ aneuploidy\ score$$

The t-value of aneuploidy coefficient $\beta_1$ was used to represent the association between RNA level and aneuploidy degree, as shown in Fig. 2F, 6E and Fig. S3C. A positive (negative) t-value represented the RNA expression of the certain gene increased (decreased) in hCEC clones with higher degree of aneuploidy.

The Log2FC of reversine/AZ3146-treated hCECs and the t-value of aneuploidy coefficient of hCEC clones were used in Fig. 5G, 6A and Fig. S5C-D. For PCA of RNA expression levels for lysosome or proteasome genes in hCEC clones shown in Fig. 5F, the raw count table of all genes obtained from RNAseq was pre-processed using the variance stabilizing transformation function in the DESeq2 package. Then 121 lysosome genes or 46 proteasome genes, as defined by the Kyoto Encyclopedia of Genes and Genomes (KEGG), were used to generate the PCA plot using the plotPCA function in the DESeq2 package. To assess the statistical significance of group separation in PCA plots, we performed permutation tests with 10,000 permutations. For each permutation, we calculated the mean Euclidean distance between groups in PC1-PC2 space as the test statistic. The p-value was determined as the fraction of permuted statistics that exceeded the observed test statistic. This analysis was implemented using custom R scripts with the vegan package (v2.6-4). The separation between diploid and aneuploid groups was considered significant when the permutation test p-value < 0.05.

**Analysis of published transcriptome data**

The RNA-sequencing reads for *TP53*WT RPE1 cells treated with DMSO, reversine, Baf A1 or MG132 were downloaded from the Gene Expression Omnibus (GSE60570). The RNA-sequencing reads for *TP53*KO RPE1 cells treated with DMSO or reversine were downloaded from GSE188644[109]. These raw sequencing reads were aligned using STAR, quantification using RSEM, and differential expression analysis using DESeq2, as described in the preceding section. The gene expression changes (Log2FC) were used to calculate the correlation in Fig. 5G and generate the heatmap in Fig. 6A and Fig. S5B-C. The raw counts table was normalized using the DESeq2, followed by Log2 transformed. The log2-transformed normalized PARP1 levels were compared in cells treated with different drugs, with the mean for the control being set to zero (Fig. 2D, Fig. 5I, Fig. S5I).

The Log2-transformed normalized PARP1, TIMELESS, or CEBPB RNA levels in HCT116 treated with DMSO and reversine were obtained from the supplementary table 7 of Cohen-Sharir et al., 2021[97], which were used in Fig. 2D, 6F and Fig. S3B, with the mean for the control being set to zero. The Log2FC of *PARP1* RNA levels in aneuploid clones generated using MMCT, compared to their near-diploid controls, were obtained from supplementary table 2 of Dürrbaum et al., 2014[30], which were used in Fig. S2I. The Log2-transformed transcript per million (TPM) of PARP1 and CEBPB in CCLE cancer cell lines were obtained from Depmap portal[110,111] (Expression public 21Q4), which were used in Fig. 4B-C, Fig. S6G, Fig. 6K. The FPKM of PARP1 in TCGA (The Cancer Genome Atlas) COREAD samples (n = 597) were obtained from the Human Protein Atlas, which were used in Fig. 4K. For CCLE cell lines, the PARP1 SCNA (Log2, obtained from Depmap portal, 21Q4) was substrate from the *PARP1* RNA levels (Log2) to normalize the copy number change. For TCGA COREAD samples, the patients with PARP1 DNA gain or loss were removed from the analysis (absolute SCNA value ≥ 0.05). To study the impact of CEBPB copy number change on PARP1 expression changes, TCGA COREAD samples without PARP1 copy alterations were classified based on CEBPB copy number status (diploid, gain, or amp, defined by cBioPortal). The PARP1 expression profile of each group is shown in Fig. S6D.

The *PARP1* RNA levels of paired and unpaired primary and metastatic tumors were downloaded from Gene Expression Omnibus (GSE131418)[52]. For the unpaired tumors, COREAD patients were separated into a discovery cohort (MCC dataset with 333 primary tumors and 184 metastases) and a validation cohort (Consortium dataset with 545 primary tumors and 72 metastases). The probe-level raw data generated via Rosetta/Merck human RSTA custom Affymetrix 2.0 microarray was read using ReadAffy function and the robust multichip average (RMA) expression was calculated using rma function in the affy package (v1.72.0)[112], including background correction, normalization, and expression calculation (Log2). Five probes for PARP1 were selected and their normalized levels in different samples were shown in Fig. S4E. The normalized levels of probe AK225654at in different samples were shown in Fig. 4N. For the paired tumors, the metastases profiles were normalized to the paired primary as processed by Zhang et al.[113]. In brief, the probe with the highest variability across patients for each gene (by SD applied to log2-transformed expression values) was selected to represent the gene in the dataset. Then the log2-transformed expression values for each metastasis expression profile were centered on its primary pair, with the values for the primary pair being set to zero. Then, the centered expression values were divided by the SD across the centered metastasis and primary profiles. The normalized expression values of PARP1 were used in Fig. 4M.

The following datasets from Gene Expression Omnibus were used to compare PARP1 or CEBPB RNA expression change in Baf A1-treated cells compared with that in DMSO-treated controls: RPE1 treated with 0.1 μM Baf A1 for 6 hours (n = 3; data from GSE60570, see above)[29], HCT116 treated with 5 nM Baf A1 for 48 hours (n = 2; data from GSE150523)[53]. HEK293T treated with 10 nM Baf A1 for 24 hours (n = 3; data from GSE141507)[54]. HeLa cells were treated with 15 nM Baf A1 for 24 hours (n = 2; data from GSE16870)[55]. The following datasets from Gene Expression Omnibus were used to compare PARP1 RNA expression change in Bortezomib- or MG132-treated cells compared with that in DMSO-treated controls: RPE1 cells treated with 1 µM MG132 for 24 hours (n = 3, data from GSE60570, see above)[29]. HCT116 treated with 5 nM bortezomib for 6 hours (n = 2, data from GSE95513)[56], 100 nM bortezomib for 3 hours, or 20 µM MG132 for 3 hours (n = 4, data from GSE165325)[58]. iPSC-derived endothelial cells (SCVI-100 and SCVI-114) or primary human aortic endothelial cells treated with 2 nM bortezomib for 72 hours (n = 1, data from GSE217898)[57]. The log2-transformed normalized PARP1 levels were directly obtained using the online analysis tool (GEO2R) of Gene Expression Omnibus, with the mean for the controls being set to zero.

**Liquid-chromatography mass spectrometry and data processing**

The cell pellets were lysed using the following buffer: 8 M urea, 100 mM Tris, pH = 8.5, 10 mM TCEP, and 40 mM CAA (150 µL per sample) and sonicated in a probe sonicator for 1 × 5 s cycle at amplitude of 50%. After sonication, the lysates were incubated for 30 min at 56°C in a thermoshaker at 1000 rpm. Insoluble debris was removed by centrifugation for 5 minutes at 16,000 × g. Protein concentrations were measured using the A280 method, and the proteins were subsequently digested with trypsin at 50:1 (w/w) ratio at 37°C (lysates were diluted sixfold with 20 mM Tris, pH = 8 prior to digestion). After digestion, the samples were acidified with 10% FA to a final concentration of 0.5% FA and centrifuged to remove undigested material. The resulting peptides were desalted using tC18 Waters SepPak cartridges, and eluates were dried on a speedvac.

50 µg of the digest from each sample were solubilized in 20 µL of 50 mM HEPES buffer pH = 8.5. Then 8 µL of 12.5 mg/mL TMTPro reagent were added, and the labeling reaction was allowed to proceed for 30 min at room temperature. Excess label was quenched by adding 40 µL of 500 mM ABC buffer for 30 min at 37°C. Labeled peptides from different samples were mixed to create two 16-plex TMT batches, which were then desalted using tC18 SepPak cartridges, concentrated using a speedvac, and fractionated offline.

500 µg of peptides were fractionated using a Waters XBridge BEH 130A C18 column (3.5 µm, 4.63 × 250 mm) on an Agilent 1260 Infinity series HPLC system operating at a flow rate of 1 mL/min with three buffer lines: buffer A consisting of water, buffer B of acetonitrile, and buffer C of 100 mM ammonium bicarbonate. Peptides were separated by a linear gradient from 5% B to 35% B in 62 min followed by a linear increase to 60% B in 5 min, and ramped to 70% B in 3 min. Buffer C was constantly introduced throughout the gradient at 10%. Fractions were collected every 60 seconds. Fractions from 30 to 64 were used for LC-MS/MS analysis.

The LC separation was performed online on EvosepOne LC (Bache et al., 2018) utilizing Dr Maisch C18 AQ analytical column (1.9 µm, 0.15 x 15 mm, Cat# EV-1106). Peptides were gradiently eluted from the column directly to an Orbitrap HFX mass spectrometer using a 44-minute evosep method (30SPD) at a flow rate of 220 nL/min. The mass spectrometer was operated in a data-dependent acquisition mode (DDA). High-resolution full MS spectra were acquired with a resolution of 120,000, an AGC target of 3e6, a maximum ion injection time of 100 ms, and scan range of 400–1600 m/z. Following each full MS scan, 20 data-dependent HCD MS/MS scans were acquired at the resolution of 60,000, AGC target of 5e5, maximum ion time of 100 ms, one microscan, 0.4 m/z isolation window, normalized collision energy of 30, fixed first mass 100 m/z, and dynamic exclusion for 45 s. Both MS and MS2 spectra were recorded in profile mode.

The MS data were analyzed using MaxQuant software (v1.6.15.0)[114] and searched against the SwissProt subset of the human UniProt database containing 20,430 entries. Database search was performed in Andromeda[115] integrated in the MaxQuant environment. A list of 248 common laboratory contaminants included in MaxQuant was also added to the database as well as reversed versions of all sequences. For searching, the enzyme specificity was set to trypsin with the maximum number of missed cleavages set to 2. The precursor mass tolerance was set to 20 ppm for the first search used for nonlinear mass recalibration and then to 6 ppm for the main search. Oxidation of methionine was searched as variable modification; carbamidomethylation of cysteines was searched as a fixed modification. TMT labeling was set to lysine residues and N-terminal amino groups, and corresponding batch-specific isotopic correction factors were accounted for. The FDR for peptide, protein, and site identification was set to 1%, and the minimum peptide length was set to 6. To transfer identifications across different runs, the 'match between runs' option in MaxQuant was disabled. Only precursors with minimum precursor ion fraction (PIF) of 75% were used for protein quantification. Match between runs option was enabled and RAW TMT reporter ion intensities of peptide features were used for subsequent data analysis in MSstatsTMT[116]. Subsequent data analysis was performed in either Perseus[117] or R for statistical computing and graphics.

The log2-transformed normalized protein levels were fitted by aneuploidy degrees of hCEC clones with a linear model as the following formula:

$$protein\ log2FC \sim \beta_0 + \beta_1 \times aneuploidy\ score$$

The t-value of aneuploidy coefficient $\beta_1$ was used to represent the association between protein level and aneuploidy degree, as shown in Fig. 2G. A positive (negative) t-value represented the protein expression of the certain gene increased (decreased) in hCEC clones with higher degree of aneuploidy.

**Analysis of published proteome data**

The proteome changes of the aneuploid clones, generated via MMCT, compared to their near-diploid controls, were obtained from supplementary table 1 of Stingele et al., 2012[32]. To compare the protein levels of these clones, stable isotope labeling with amino acids in cell culture (SILAC) was used, followed by high-resolution mass spectrometry and quantified the proteome to a depth of ~6000 proteins. This data was used in Fig. 2I and Fig. S2I. The proteome data of CCLE cancer cell lines were obtained from Depmap portal[110,111] (Expression public 21Q4), which was used in Fig. 4B-C.

##### Resistance of cancer cell lines to olaparib

The data of cancer cell sensitivity to the PARP1 inhibitor Olaparib were obtained from Depmap portal. The following three independent drug screening datasets were used: PRISM repurposing primary screen[118] (olaparib ID: BRD-BRD-K02113016-001-15-4, n = 547 cell lines), GDSC dataset[119] (olaparib ID: GDSC21017, n = 506 cell lines) and CTD2 dataset[120] (olaparib ID: CTRP411867, n = 756 cell lines). For the PRISM dataset, the drug effect log2FC data was downloaded, while the AUC values of olaparib in the GDSC and CTD2 datasets were used. The Z-values of these values (Log2FC or ACU) in each dataset were calculated and used to generate the box plot in Fig. S6I. A higher Z-value represents an increased level of resistance to olaparib.

##### Patients' survival analysis using TCGA

The TCGA survival data of COREAD patients was downloaded from the Human Protein Atlas. Based on the FPKM value of PARP1, patients were classified into two expression groups (PARP1-Low and PARP1-High) to yield maximal difference with regard to survival between the two groups at the lowest log-rank p-value (n = 124 patients with low PARP1 and n = 473 patients with high PARP1). The prognosis of each group of patients was examined by Kaplan-Meier survival estimators, and the survival outcomes of the two groups were compared by log-rank tests.

##### Knockdown, knockout, and overexpression

The PARP1 overexpression and *PARP1*, *GPX1*, *CAT*, *TFEB*, *TFE3* knockdown plasmids, along with their respective controls, were used to generate lentivirus with packaging and envelope plasmids. hCECs and COREAD cancer cells were transduced with the lentivirus, followed by selection using puromycin (1.5 μg/mL for hCECs and 3 μg/mL for COREAD cancer cells with shRNA) or blastomycin (10 μg/ml for hCECs and COREAD cancer cells with pHAGE-CMV-blast) until the uninfected control cells died. For the co-expression of the shRNA targeting the 3'-UTR region of *PARP1* transcripts and shRNA-resistant PARP1 cDNA, the PARP1-overexpressing cells were transduced with shPARP1-1, followed by puromycin selection. To get rid of unspecific impacts of lentiviral infection and selection, all cells in Fig. 3G, including those lacking shPARP1-1 or *PARP1* OE, and those exclusively expressing shPARP1-1 or *PARP1* OE, were additionally infected with the control virus for a total of two rounds of infection and selection. For example, the cells without shPARP1-1 or *PARP1* OE were subsequently infected and selected for pHAGE-EV and shNC. The effects of PARP1 overexpression and knockdown were confirmed using WB. To achieve physiological levels of PARP1 expression, *TP53*WT or *TP53*KO hCECs were transduced with a lentiviral Tet-On system for doxycycline-inducible PARP1. Following hygromycin selection, cells were treated with 1 μg/mL doxycycline for 1-2 days to induce PARP1 expression before validation by WB or assessment of H2O2 sensitivity. To establish stable co-expression of *PARP1* and *TIMELESS* in hCEC A29 clones, cells were first transduced with pHAGE-PARP1-blast or pHAGE-EV-blast lentivirus and selected with blastomycin. The stable cells were then transduced with pHAGE-TIMELESS-hygro or pHAGE-EV-hygro lentivirus, followed by hygromycin selection. For CEBPB overexpression, cells were transduced with lentivirus generated from pBABE-puro LAP2 (Addgene #15712) or pBABE-puro empty vector. For *NRF2*, *PARP1*, *PARP2*, and *MIF* knockout, lentiviral particles were produced by co-transfecting HEK293T cells with LentiCRISPR-V2-FE-puro plasmids carrying target-specific sgRNAs or non-targeting control (sgNC), along with lentiviral packaging and envelope plasmids. Target cells were transduced with the lentivirus and selected with puromycin. For *PARP1* knockout and *TIMELESS* expression in hCEC A29 clones, cells were co-transduced with pHAGE-TIMELESS-hygro (or pHAGE-EV-hygro) and LentiCRISPR-V2-FE-puro plasmid carrying PARP1 sgRNA (or sgNC) lentivirus, followed by hygromycin and puromycin selection before validation by WB or assessment of H2O2 sensitivity.

**Cancer cells expressing luciferase and GFP or mCherry**

For the comparison of ROS sensitivity of near-diploid and high-aneuploidy colon cancer cell lines using the multicolor competition assay, as well as the observation of intravenously injected cancer cells in living mice (see below), we lentivirally transduced two near-diploid colon cancer cells (DLD1 and HCT116) and one high-aneuploid colon cancer cells (HT29) with either pCDH-EF1a-eFFly-eGFP (Addgene, 104834) or pCDH-EF1a-eFFly-mCherry (Addgene, 104833). After infection, the GFP-positive or mCherry-positive cells were sorted using flow cytometry-based cell sorting (FACS) on the SONY SH800 cell sorter with a 130 μm sorting chip.

**Multicolor competition assay**

To investigate the impact of *PARP1* knockdown or overexpression on the sensitivity of colon cell lines to $H_2O_2$, control cancer cells, labeled with either GFP or mCherry, were mixed with PARP1-knockdown or PARP1-overexpressing cells labeled with the complementary fluorescent marker (i.e., mCherry or GFP) in equal cell numbers. The six combinations examined were as follows: HCT116-GFP with shNC and HCT116-mCherry with shPARP1-1, HCT116-mCherry with shNC and HCT116-GFP with shPARP1-1, DLD1-GFP with shNC and DLD1-mCherry with shPARP1-1, DLD1-mCherry with shNC and DLD1-GFP with shPARP1-1, HT29-GFP with pHAGE-EV and HT29-mCherry with pHAGE-PARP1, HT29-mCherry with pHAGE-EV and HT29-mCherry with pHAGE-PARP1. The mixed cells were cultured in the regular medium, with and without the addition of 2mM $H_2O_2$, for a 24-hour incubation period. Then the ratios of GFP-labeled and mCherry-labeled cells within each combination were quantified using flow cytometry (SONY, SA3800). To assess the impact of $H_2O_2$ treatment, these ratios were normalized to those obtained from control groups that were not exposed to $H_2O_2$.

**Assessing metastatic potential through in vivo assay**

The animal procedures were approved by the New York University Grossman School of Medicine's Institutional Animal Care and Use Committee (IACUC). All the procedures are conducted in facilities approved by the Association for Assessment and Accreditation of Laboratory Animal Care International (AAALAC).

To assess dissemination potential, luciferase-expressing human colon cancer cells (DLD1 and HCT116 with shNC or shPARP1-1, HT29-EV, and SW498) were intravenously injected into immunodeficient NOD.Cg-Prkdc$^{scid}$ Il2rg$^{tm1Wjl}$/SzJ (NSG) mice. For immune-competent studies, luciferase-expressing aneuploid mouse KP lung cancer cells with empty expression or PARP1 overexpression vectors were injected into B6 Albino mice. Tumor progression was monitored by bioluminescence imaging every two weeks until the pre-established endpoint.

For the intravenous injection, cancer cells were collected through trypsinization, followed by two washes with cold PBS and suspension in cold PBS at a concentration of 1,000,000 cells/mL. Then 200,000 cancer cells in 200 µL PBS were intravenously injected (via tail vein) into female NSG or B6 Albino mice aged 7-9 weeks, using a 1 mL syringe with a 30G, 0.3 x 13mm needle. Each type of cancer cell was introduced into at least five mice. Continuous monitoring of the mice's condition was performed throughout the procedures and up to 1-2 hours after the injection.

For in vivo bioluminescence imaging, the mice were anesthetized via inhalation with the oxygen level regulator set between 1-2 L/min and the isoflurane vaporizer set at 2.5-5% for induction and 1.5–3% for maintenance. The D-luciferin substrate (150 mg/kg) was administered to mice via intravitreal injection using a 0.5 mL insulin syringe, 28 G needle. Then the imaging was performed using a bioluminescence imaging (BLI) scanner (IVIS Lumina XRMS In Vivo Imaging System). For data analysis, regions of positive signal were defined for each mouse while carefully excluding the ears and nose. The total luminescent flux (p/sec) within the regions of positive signal was quantified and normalized to the initial levels immediately following intravenous injection. For mice which reached the pre-established endpoint in the middle of the observation window (10 weeks), they are labeled by crosses in the figure and their last total flux signals were used for statistical analysis for the later time points.

To compare tumor growth between aneuploidy states in immune-competent environments, KP lung cancer cells with high or low-aneuploidy were harvested at subconfluency, washed with PBS, and resuspended at $1 \times 10^6$ cells/mL. Each mouse received $2 \times 10^5$ cells via tail vein injection. Tumor volumes were quantified using MRI. For MRI quantification, mice were anesthetized with isoflurane, and lung fields were scanned using a BioSpec USR70/30 horizontal bore system (Bruker) to obtain 16 consecutive sections. MRI signal acquisition was gated to cardiac and respiratory cycles to minimize motion artifacts. Whole lung tumor volumes were quantified from volumetric measurements using 3D Slicer software.

##### Fluorescence-activated cell sorting based on PARP1 levels

To sort cells based on their PARP1 expression levels, 60,000,000 cells were collected through trypsinization, followed by a wash with cold PBS and fixation with 10 mL of 2% paraformaldehyde solution in PBS for 7 minutes at room temperature. After a wash with cold PBS, the cells were suspended in 10 mL of 90% cold methanol and incubated for 10 minutes at 4℃. Then the cells were washed with 10 mL of cold 1% BSA in PBS and blocked with 10 mL of 3% BSA in PBS for 45 minutes at 4°C in the dark. Subsequently, the cells were incubated with 6 mL of CoraLite 488-conjugated PARP1 monoclonal antibody (Proteintech, CL488-66520) at a concentration of 2 μg/mL in 1% BSA in PBS for 60 minutes at 4°C in the dark. Then the cells were washed three times with cold 1% BSA in PBS and resuspended in cold 1% BSA in PBS at a concentration of 5,000,000 cells/mL for sorting.

For the fluorescence-activated cell sorting, the SONY SH800 cell sorter with a 100 μm sorting chip was used. After excluding debris (gate 1: X-axis: FSC-A, Y-axis: BSC-A) and doublets (gate 2: X-axis: FSC-A, Y-axis: FSC-H), two gates above and below the diagonal line of the main populations on the plot of PARP1 intensity (FITC-A) versus cell size (BSC-A), representing PARP1-low and PARP1-high populations, respectively, were chosen, with each selected population comprising ~10% of the total cell count. After collection of the PARP1-low and PARP1-high cells, they were centrifuged to remove the supernatant. The cell pellets were stored at -80°C.

##### Genome-wide CRISPR screen

To unbiasedly identify genes that modulate PARP1 expression, we conducted two independent, fluorescence-activated cell sorting (FACS)-based genome-wide CRISPR screens. The cellular level of PARP1 served as the readout in these screens. The sensitivity of PARP1 antibody was validated using PARP1-overexpressed cells, PARP1-knockdown cells, and hCEC clones with known different levels of PARP1 expression. For each screen replicate, 60,000,000 hCEC cells were infected with a library of lentiviruses harboring 94,335 sgRNAs targeting 19,026 protein-coding human genes at a multiplicity of infection (MOI) lower than 1 (maintaining a representation ~450 per gRNA). After puromycin selection and a brief expansion, the cells were divided into two or three technical replicates (60,000,000 cells for each technical replicates), fixed and immunostained with the fluorophore-conjugated PARP1 antibody, and sorted based on PARP1 protein expression levels (see above). The sorting process resulted in approximately 4 million cells per group (two groups: PARP1-high or PARP1-low) in each technical replicate (two or three technical replicates for each biological replicate, two biological replicates in total), accounting for approximately 10% of all cells. To ensure that the FACS-based readout of PARP1 level was highly specific and sensitive, we validated the antibody using PARP1-overexpressed cells, PARP1-knockdown cells, and hCEC clones with known different levels of PARP1 expression (Fig. 5B).

The genomic DNA was then extracted from the sorted cells with low or high PARP1, and the unsorted cells as control. The following procedure shared by Neville Sanjana Lab was used for genomic DNA extraction from 40,000,000 cells. For different amounts of cells, the quantities were scaled proportionally. In a 15 mL conical tube, 6 mL of NK Lysis Buffer (50 mM Tris, 50 mM EDTA, 1% SDS, pH 8) and 30 μL of 20 mg/mL Proteinase K (Qiagen, 19131) were added to the cell sample and incubated at 55 °C for 24 hours. Then 30 μL of 10 mg/mL RNAse A (Qiagen, 19101) was added to the lysed sample, which was then inverted 25 times and incubated at 37 °C for 30 minutes. Samples were cooled on ice before addition of 2 mL of pre-chilled 7.5 M ammonium acetate (Sigma-Aldrich, A1542) to precipitate proteins. Subsequently, the samples were vortexed at high speed for 20 seconds and then centrifuged at ≥ 4,000 x g for 10 minutes. After the spin, a tight pellet was visible in each tube and the supernatant was carefully decanted into a new 15 mL conical tube. Then 6 mL of 100% isopropanol was added to the tube, inverted 50 times and centrifuged at ≥4,000 x g for 10 minutes. Genomic DNA was visible as a small white pellet in each tube. The supernatant was discarded, 6 mL of freshly prepared 70% ethanol was added, the tube was inverted 10 times, and then centrifuged at ≥4,000 x g for 1 minute. The supernatant was discarded by pouring; the tube was briefly spun, and remaining ethanol was removed using a P200 pipette. After air drying for 10-30 minutes, the DNA changed appearance from a milky white pellet to slightly translucent. At this stage, 500 μL of TE buffer was added, the tube was incubated at 65 °C for 1 hour and at room temperature overnight to fully resuspend the DNA. The next day, the gDNA samples were vortexed briefly. The gDNA concentration was measured using a Nanodrop (Thermo Scientific).

The DNA of sgRNAs was extracted from the genomic DNA using a 26-cycle PCR reaction. Each PCR reaction consisted of 4 μg genomic DNA, 0.5 μM primers and 1 unit of Q5 DNA polymerase (NEB, M0491L) in a 100 μL PCR reaction volume. All extracted genomic DNA from the sorted cells and the same amount of genomic DNA from the unsorted cells were used for this round of PCR. Then all the PCR products from the same sample were pooled together, and 100 μL of PCR product for each sample was gel purified and eluted in 35 μL of the elution buffer. The concentration of the purified PCR product was determined using the Qubit dsDNA quantification assay kit (Thermo Scientific, Q32854), and 400 ng of the purified PCR products were used in a 5-cycle PCR reaction to add adaptors. Next, 5 μL of PCR product for each sample was used in a 7-cycle PCR reaction to introduce the standard Illumina sequencing primers and unique sample barcodes. The concentrations of PCR products were determined by analyzing the band intensities after gel electrophoresis. Equimolar PCR products from the sorted samples and two-fold PCR products from the control samples were mixed together, followed by the gel electrophoresis purification using the Nucleospin Gel and PCR Clean-UP kit. The sequence of all the primers is included in Table 7.

The prepared library was sequenced on an Illumina NextSeq 500 in single-end mode (1 X 75 cycles). Approximately 10 million reads were obtained for each sorted sample and about 20 million reads were obtained for each control sample. The raw reads were trimmed using cutadapt [121] with the adaptor sequence ACGAAACACC. Subsequently, the trimmed reads were processed to generate a count table using the count function of the MAGeCK package[122] and calculate the enrichment scores (MAGeCK values) at the gene level using the mle function of the MAGeCK package. The sgRNAs targeting positive regulators of PARP1 expression (PARP1 activators) were enriched in the PARP1-low cells compared to the unsorted cells, while sgRNAs targeting negative regulators of PARP1 expression (PARP1 suppressors) were enriched in the PARP1-high cells. To plot the relation of MAGeCK scores in the PARP1-low and these in the PARP1-high population (Fig. 5C), genes exclusively enriched in the PARP1-high population (the 2nd quadrant) and in the PARP1-low population (the 4th quadrant) are depicted in red and green, respectively. The intensity of the dots represents gene-level FDR-adjusted p-values within their respective populations (-Log10). For genes located in the other two quadrants, their p-values were set to one.

**Enrichment analysis using Ingenuity Pathway Analysis (IPA)**

The CRISPR screen identified 269 PARP1 activators (genes whose knockout decreased PARP1 expression) and 155 PARP1 suppressors (genes whose knockout increased PARP1 expression) using a false discovery rate (FDR) cutoff of $< 0.1$ and a MAGeCK beta value cutoff of $> 0.5$. The two sets of the filtered genes were used as the input for the pathway analysis performed using QIAGEN IPA (Expression analysis of the IPA core analysis with the measurement type being expr other). Enriched pathways with B-H p-values smaller than 0.1 are shown.

To identify the genes whose expression change in aneuploid cells may contribute to PARP1 downregulation, we searched for the PARP1 activators downregulated in aneuploid cells and the PARP1 suppressors upregulated in aneuploid cells (referred to as aneuploid-associated PARP1 regulators). For the MPS1i-induced aneuploid cells, RNA log2FC was used, while the t-values of aneuploidy coefficient was used for hCEC clones (see above). Genes with an FDR larger than 0.1 or the absolute Z-scores $\leq 0.5$ were considered unchanged. Only the genes that exhibited expected expression changes in at least two aneuploidy models, without conflicted changes in different models were considered as aneuploid-associated PARP1 regulators, which were used as the input for the pathway analysis as described above.

**Identification of potential PARP1 regulators contributing to its downregulation in aneuploid cells**

Decreased PARP1 RNA in aneuploid cells and cells with lysosomal dysfunction suggested altered PARP1 transcription. We aimed to identify transcription factors (TFs) responsible for PARP1 transcriptional dysregulation in these settings. We integrated our PARP1 CRISPR screen results with TF databases[59–61] to identify PARP1-regulating TFs. We used three published databases: the GeneHancer database from GeneCards, TFLink, and AnimalTFDB[59–61]. All potential PARP1 TFs from these databases were merged for further analysis.

The TFs were filtered based on our PARP1 screen results into two groups: TFs promoting PARP1 expression (promoters, enriched in PARP1-low cells or depleted in PARP1-high cells, $p<0.05$) and TFs suppressing PARP1 expression (suppressors, depleted in PARP1-low cells or enriched in PARP1-high cells, $p<0.05$). We then examined TF expression changes in aneuploid and lysosome-dysfunction cells, identifying downregulated promoters and upregulated suppressors ($p<0.05$). This analysis identified CEBPB as a potential PARP1 suppressor, showing consistent upregulation across aneuploid and lysosome-dysfunction models.

**ENCODE CEBPB ChIP-Seq data**

The CEBPB ChIP-Seq data (bigWig files for signal p-values and bigBed narrowPeak files for IDR thresholded peaks) for A549 cells treated with or without CEBPB agonist dexamethasone (1 hour) were downloaded from ENCODE database (ENCSR625DZB). The signal p-values and IDR thresholded peaks near PARP1 TSS were visualized using IGV software (V2.17.4).

**ChIP Quantitative PCR**

To assess CEBPB binding to the *PARP1* promoter in different conditions, we performed chromatin immunoprecipitation followed by quantitative PCR (ChIP-qPCR). Dexamethasone (100nM) treated cells were used as a positive control. A total of 10,000,000 cells were harvested by trypsinization. The cells were first double-crosslinked with 1.5 mM EGS (ethylene glycol bis(succinimidyl succinate) for 10 minutes at room temperature, followed by 4% formaldehyde for an additional 30 minutes. Crosslinking was stopped by treating the samples with 125 mM glycine at room temperature with gentle rocking for 5 minutes. The crosslinked cells were washed with PBS + 0.5% BSA and then resuspended in 1200 µLL ChIP lysis buffer (10 mM Tris-Cl, pH8, 100 mM NaCl, 1 mM EDTA, 0.5 mM EGTA, 0.1% Sodium deoxycholate, 0.5% N-lauroylsarcosine and 1% Triton X-100) supplemented with protease inhibitor cocktail and incubated at 4$^{o}$C on a revolver for 10 minutes. The chromatin was then sheared using a Diagenode Bioruptor Pico sonication device at ultra-high intensity for 300 cycles of 45 seconds ON and 30 seconds OFF. To ensure that the size of sheared chromatin was between 200-500 bp, 5 µL of the sheared chromatin was reverse crosslinked by incubating with 85 µL of 10mM TrisCl, pH 8.0 and 2 µL of RNase A (10 mg/mL) at 37$^{o}$C for 30 minutes followed by addition of proteinase K (20 mg/mL) and incubation at 55$^{o}$C for 2 hours on a thermomixer at 800 rpm. The reverse crosslinked chromatin was purified using AMPure DNA purification beads and the fragment size was analyzed using Agilent 4150 TapeStation System. Lysates were used as the input reference control. For pulldown, 50 µL protein A dynabeads were washed twice with PBS-BSA and then resuspended in 400 µL PBS-BSA with 5 µg of antibody against either CEBP beta (Abcam, ab32358) or IgG control (Cell Signaling, 3900) and incubated for 1 hour at room temperature on a revolver. After incubation, the antibody conjugated beads were removed from the suspension and resuspended in 100 µL of ChIP lysis buffer and added to the sonicated chromatin samples and incubated overnight at 4°C. Next day, the beads with immunoprecipitated samples were washed thrice with wash buffer I (20 mM Tris-Cl, pH 7.5, 150 mM NaCl, 2 mM EDTA, 0.1% SDS and 1% triton X-100), twice with wash buffer II (10 mM Tris-Cl, pH 8, 120 mM LiCl, 1mM EDTA, 0.7% sodium deoxycholate and 1% triton X-100), twice with TET buffer (10 mM Tris-Cl, pH 8, 1 mM EDTA, 0.2% Tween-20) and then eluted in 70 µL elution buffer (10 mM Tris-Cl pH 8, 300 mM NaCl, 5 mM EDTA and 0.5% SDS). The pure chromatin was then reverse crosslinked and purified as mentioned above. Concentration of the purified DNA fragments was measured using Qubit 2.0 fluorometer and the Qubit dsDNA HS kit and their size was assessed on the tapestation system. Next, the enrichment of the ChIP and input DNA was quantified and analyzed by qPCR using two independent primer sets within the PARP1 promoter: set of primer 1 (Forward 5'GATTGTTCTGTCCCAGGAAGT3', Reverse 5'GACTGCAGTGAGCCATGAT3'), set of primer 2 (Forward 5'GCTAGTAGCTCTTTGGAGGAC3', Reverse 5'CTCAGGAGTTCCAGACTGC3'), and a negative control primer set (Forward 5'TCCACCCACTTCTCTCCATCT3', Reverse 5'GCTGCAAGTAATGGGCTCAG3'). The resulting ChIP-qPCR data was analyzed by using the percent input method and normalized to negative control.

**Analysis of published scRNA-Seq and CRISPRa Perturb-Seq data**

The raw unique molecular identifier (UMI) counts, cellular annotations, and inferred copy number alterations (CNVs) for GSE178341 were obtained from the Curated Cancer Cell Atlas (https://www.weizmann.ac.il/sites/3CA/)[66,67]. To ensure high-quality scRNA-seq data, we applied rigorous quality control (QC) criteria: first, cells with fewer than 500 or more than 8000 detected genes were removed to eliminate low-quality and potential multiplet cells. Second, cells with mitochondrial gene content exceeding 20% were excluded, as high mitochondrial RNA expression is often indicative of apoptotic or stressed cells. Third, malignant cells were subset for downstream analysis to focus on tumor-specific transcriptomic and genomic alterations.

Data preprocessing was conducted using Seurat v4 in R[123]. SCTransform v1 was applied to normalize raw UMI counts[124]. Harmony was used to correct batch effects across samples, ensuring effective dataset integration while maintaining cellular identity[125]. The average inferred CNV signal across chromosomes was computed to estimate chromosomal instability. Aneuploidy Score (AS) was calculated based on the defined formula for quantifying genomic instability[12]. To investigate the functional relevance of lysosomal activity in malignant cells: the KEGG lysosome gene set (MSigDB, M11266) was used to compute the Seurat module score.

Due to the zero-inflation characteristic of scRNA-seq data, additional filtering was applied before performing correlation analysis and UMAP Visualization: Cells with a combined expression of PARP1 and CEBPB below 4 were excluded from correlation and UMAP analyses to improve signal reliability. In the correlation analysis, log1p transformation is applied on SCTransform corrected UMI counts of CEBPB and PARP1, and Pearson correlation is applied to reveal the negative correlation. UMAP projections were aestheticized using Kernel Density Estimation (KDE), showing cells above the threshold on gene expression, lysosome module score (top 30% of all malignant cells) and aneuploidy score (>5), with density weights based on gene expression, aneuploidy score, and lysosome module score, improving interpretability in visualizations.

The CRISPRa Perturb-Seq data relevant to cells with CEBPB activation and non-targeting control was provided by Dr. Thomas Norman. The CEBPB and PARP1 read counts after total UMI count normalization were used in Fig. 6J (left panel). The PARP1 expression z-scores relative to expression levels in control cells were used in Fig. 6J (right panel).

**Replicates and statistical analyses**

In all figures, individual data points represent replicates from representative experiments (at least two independent experiments were performed to validate results). For dot plots, unless otherwise specified, summary statistics show mean ± standard deviation. All statistical analyses are two-sided unless otherwise specified, with statistical significance indicated as: ns, $p>0.05$; *$p<0.05$; **$p<0.01$; ***$p<0.001$; ****$p<0.0001$. The test statistics with confidence intervals, effect sizes, degrees of freedom and P value are detailed in Table 7.

**Material Resource**

The source and identifier for all materials used in this study are listed in Table 7 and STAR Methods.

**Code availability**

The code used to generate and/or analyze the data is available upon request.

**Data availability**

All datasets are available within the article and its supplementary information, or from the corresponding Author upon request.

#### Supplementary Figure Legends

**Figure S1. Aneuploidy promotes tumor progression and enhances resistance to oxidative stress**
(**A**) Chromosome arm-level SCNAs in TP53KO hCECs treated with MPS1 inhibitors (reversine or AZ3146) inferred by single-cell RNA sequencing. (**B**) Viability of diploid (DMSO, GFP-labeled) versus aneuploid (AZ3146, unlabeled) TP53WT hCECs after 24h exposure to various stressors. Mean ± SD (n=4). One-way ANOVA with Dunnett's test. (**C**) Left: schematic of MPS1i treatment in confluent versus non-confluent cells to isolate off-target effects. Middle: representative fluorescent images after $H_2O_2$ treatment. Blue: Hoechst 33342 (all cells); red: PI (dead cells). Right: viability quantification. Mean ± SD of 9 fields from representative experiment of ≥2. One-way ANOVA with Dunnett's test. (**D-E**) Viability of diploid (DMSO) and aneuploid (reversine or AZ3146) (D) RPE1 (TP53KO), hPNE, hMEC, DLD1 cells, or (E) mouse KP lung cancer cells after $H_2O_2$ treatment (1 mM, 500 μM, 3.2 mM, 750 μM, and 500 μM, respectively, 24h). Mean ± SD of 3-4 replicates from representative experiment of ≥2. One-way ANOVA with Dunnett's test. (**F**) Viability after $H_2O_2$ treatment in diploid (DMSO) and aneuploid (reversine or AZ3146) TP53KO hCECs, with or without arrested cell cycle (0.5 μM palbociclib or 2 mM thymidine, 24h). Mean ± SD of triplicates from representative experiment of ≥2. One-way ANOVA with Tukey's test. (**G**) Chromosome-level SCNA in representative isogenic near-diploid (D23 and D29) and aneuploid (A20, A21, A26, and A29) hCEC clones. (**H**) Relative viability of co-cultured GFP-labeled near-diploid and unlabeled aneuploid hCEC clones (3 clones for each group) under low $H_2O_2$. Mean ± SD (n=4). One-way ANOVA with Dunnett's test (**I**) Colony formation by near-diploid and aneuploid hCECs in soft agarose gel after 10 days of growth either with or without low $H_2O_2$ (10 μM). (**J-K**) Viability after $H_2O_2$ treatment in near-diploid and aneuploid clones with (H) unattachment or (I) attachment on plastic/Geltrex surfaces. Mean ± SD. n ≥ 2. One-way ANOVA with Tukey's test. (**L**) Top: chromosome number quantification in mouse KP lung cancer cells after induction of aneuploidy (AurkB) versus control. Bottom: representative metaphase spread images. (**M**) Representative images of near-diploid (HCT116 and DLD1) and high-aneuploidy (HT29, SW403, SW948, and SK-CO-1) colon cancer cell lines without $H_2O_2$ treatment. (**N**) Viability of near-diploid and aneuploid hCEC clones after treatment with 100 μM xanthine and 20 mU/mL xanthine oxidase for 24 hours. Mean ± SD of 9 fields from representative experiment of ≥2. One-way ANOVA with Tukey's test. ns, p>0.05; *p<0.05; **p<0.01; ***p<0.001; ****p<0.0001. Scale bar, 50 μm.

**Figure S2. PARP1 is suppressed in aneuploid cells**

(**A**) Assessment of NRF2 nuclear translocation in hCECs with acute and chronic aneuploidy. Left: representative immunofluorescence images of NRF2 (green) and DAPI (blue) staining in control (untreated) or NRF2-activated cells (50 µM $H_2O_2$, 1h, or 10 µM MG132, 4h). Scale bar, 10 µm. Middle and right: quantification of nuclear/cytoplasm NRF2 ratio in diploid and aneuploid hCECs. Mean ± SD, n=25-75 cells per condition. One-way ANOVA test with Tukey's test. (**B**) Viability after $H_2O_2$ treatment in diploid and aneuploid hCECs with *NRF2* knockout. Mean ± SD of 9 fields from representative experiment of ≥3. One-way ANOVA with Dunnett's test. (**C**) Viability after $H_2O_2$ treatment in diploid and aneuploid hCEC clones with GPX1 or CAT knockdown. Mean ± SD of 4 fields from representative experiment of ≥3. One-way ANOVA. (**D**) DNA damage quantification in hCEC cells under different $H_2O_2$ concentrations using alkaline comet assay. Medians with 95% CI (n≥2). Kruskal-Wallis one-way ANOVA test with Dunnett's test. (**E**) Lipid peroxidation analysis. Left: representative flow cytometry histogram C11-BODIPY 591/510 ratio after $H_2O_2$ treatment (50 µM, 30 minutes). Middle and right: fold change quantification of 591/510 ratio in hCECs with acute or chronic aneuploidy versus their diploid controls. Mean ± SD (n=4-6). One-way ANOVA test with Dunnett's and Tukey's test for acute and chronic aneuploidy, respectively. (**F**) Viability after $H_2O_2$ treatment in hMEC cells with or without PARP1 inhibitor treatment (10 µM olaparib). Mean ± SD. Unpaired t-test. (**G**) Representative western blot of pADPr levels in diploid (DMSO) and aneuploid (reversine or AZ3146) hCECs after exposure to 50 µM $H_2O_2$ for 10 minutes. (**H**) Representative western blot of PARP1 protein levels in diploid and aneuploid TP53WT hCECs in the presence of 10 µM Z-VAD-FMK (a pan-caspase inhibitor) during DMSO or MPS1i treatment and the subsequent release. (**I**) The changes of *PARP1* DNA copy number, mRNA, and protein levels in HCT116 and RPE1 aneuploid clones (generated by MMCT) compared to the near-diploid controls. RNA expression data from Dürrbaum et al., 2014; DNA copy number and protein expression data from Stingele et al., 2012. (**J**) Representative western blot of PARP1 protein levels in near-diploid and aneuploid clones of RPE1 and HCT116 cell lines (generated via MMCT). (**K**) Representative western blot of PARP1 protein levels in near-diploid and aneuploid mouse KP lung cancer cells. ns, p>0.05; *p<0.05; **p<0.01; ***p<0.001; ****p<0.0001.

**Figure S3. Cell death and DNA repair mediated by PARP1 are suppressed in aneuploid cells by PARP1 inhibition**

(**A**) Viability of control (empty vector) and PARP1-overexpressed hCEC clones after low $H_2O_2$ treatment (20 μM). Mean ± SD of 4 fields from representative experiment of ≥2. One-way ANOVA with Tukey's test. (**B**) Relative RNA levels of *TIMELESS* (Log2) between aneuploid (reversine or AZ3146) and diploid (DMSO) hCEC, RPE1, TP53KO RPE1, and HCT116 cells. Mean ± SD, n=2-24. One-way ANOVA with Dunnett's test (hCEC); unpaired t-test (RPE1, TP53KO RPE1, and HCT116). (**C**) Correlation between aneuploidy degree and TIMELESS RNA (top) or protein levels (bottom) in hCEC clones. Linear regression is included to fit the data. (**D**) Viability of A29 clone exposed to non-lethal $H_2O_2$ (20 μM) with *PARP1* overexpression, *TIMELESS* overexpression, or both. Mean ± SD from 9 fields/replicate, n=2 replicates from representative experiment. One-way ANOVA with Tukey's test. (**E**-**F**) Viability after $H_2O_2$ treatment in near-diploid hCECs with (E) *PARP1* or *PARP2* knockout or (F) *MIF* knockout. Mean ± SD of 9 fields from representative experiment of ≥2. One-way ANOVA with Dunnett's test. (**G**) Schematic representation of the method to assess DNA damage dynamics in diploid and aneuploid hCEC clones before and after exposure to genotoxic agents. DNA damage marker γH2AX levels were measured at control, release (R0: 0h post $H_2O_2$ exposure), and recovery (R6 or R24: 6 or 24h post $H_2O_2$ exposure) timepoints. (**H**) Violin plots showing γH2AX intensity dynamics before and after treatment with various genotoxic agents in near-diploid (blue) and high-aneuploidy (red) hCEC clones. Plots show median and quartiles from 343 to 2825 cells. (**I**) Quantification of DNA damage levels (γH2AX intensity) after release from genotoxic agents compared to immediate release timepoint. Medians with 95% CI from 343 to 2825 cells. Kruskal-Wallis one-way ANOVA test with Dunnett's test. (**J**) Representative western blot of γH2AX levels 6 hours after H2O2 exposure (50 μM, 1h) in the acute aneuploidy hCEC models (TP53KO) with or without *PARP1* overexpression. (**K**) Representative western blot of γH2AX and total H2AX levels in near-diploid and aneuploid hCEC clones before and after $H_2O_2$ exposure. (**L**) Western blot of PARP1 and TIMELESS protein expression (left) and quantification of $H_2O_2$-induced (50 μM) cell viability changes (right) in A29 with *PARP1* KO, *TIMELESS* overexpression, or both. Mean ± SD from 9 fields/replicate, n=2 replicates from representative experiment. One-way ANOVA with Dunnett's test. ns, $p>0.05$; *$p<0.05$; ***$p<0.001$; ****$p<0.0001$.

**Figure S4. PARP1 downregulation promotes metastasis**

(**A**) Relative viability of HT29-GFP-EV to HT29-mCherry-PARP1 when co-cultured together, with and without 2 mM $H_2O_2$ treatment for 24 hours. Mean ± SD of 4 replicates from representative experiment of ≥2. Unpaired t-test. (**B**) The relative cell number ratio of DLD1-GFP-shNC (or HCT116-GFP-shNC) to DLD1-mCherry-shPARP1 (or HCT116-mCherry-shPARP1) when co-cultured together, with and without 2 mM $H_2O_2$ treatment for 24 hours. Mean ± SD of 3 replicates from representative experiment of ≥2. Unpaired t-test. (**C**) Doubling time comparison of DLD1 and HCT116 cells with or without *PARP1* knockdown, and HT29 cells. Mean and SD from 9 fields/replicate, n=12 replicates from representative experiment. One-way ANOVA with Tukey's test. (**D**) Doubling time comparison of mouse KP lung cancer cells with empty vector (EV) and mouse *PARP1* (mPARP1) overexpression. Mean ± SD from 9 fields/replicate, n=8 replicates from representative experiment. Unpaired t-test. (**E**) PARP1 RNA levels in primary versus metastatic COREAD tumors across different PARP1 probes (Rosetta/Merck human RSTA custom Affymetrix 2.0 microarray) in discovery (top) and validation (bottom) datasets. Boxplots show median with first/third quartiles. Unpaired t-test. ns, p>0.05; *p<0.05; **p<0.01; ***p<0.001; ****p<0.0001.

**Figure S5. Genome-wide CRISPR screen reveals lysosomal stress as a potential mediator of PARP1 downregulation in aneuploid cells**

(**A**) The gating strategy used to sort PARP1-low (gate P) and PARP1-high (gate R) populations during fluorescence-activated cell sorting (FACS) for the genome-wide CRISPR screens (**B**) Pathway analysis of 155 *PARP1* suppressors identified by CRISPR screen (enriched in PARP1-high cells, beta value > 0.5 and FDR < 0.1). Pathways with adjusted p-values < 0.1 are shown. (**C**) Heatmap showing expression changes of identified *PARP1* activators, suppressors, and predicted upstream factors in aneuploid cells compared to diploid cells. Expression changes are shown as Z-scores of RNA log2FC (reversine- or AZ-treated cells) or t-values (hCEC clones). Genes with FDR > 0.1 were set to a Z-score of zero. The expected expression change is only directional (increase or decrease) and predicted to result in PARP1 downregulation. (**D**) Left: workflow for identifying PARP1 regulators that potentially contribute to PARP1 downregulation in aneuploid cells. Expression changes were filtered using a Z-score threshold of ±0.5. Genes showing consistent predicted expression changes across multiple aneuploidy models (n≥2) were identified as potential PARP1 regulators. Right: heatmap showing expression changes of the identified PARP1 regulators. (**E**) Pathway analysis of PARP1 regulators identified in (C) (n=65 genes). Pathways with adjusted p-values < 0.01 are shown. Blue bars indicate predicted pathway inhibition. (**F**) Rank plots showing enrichment and depletion of sgRNAs in PARP1-low (top) and PARP1-high (bottom) sorted populations relative to unsorted samples from the CRISPR screen. Blue and red indicate depleted and enriched sgRNAs respectively. (**G**) Spearman correlation of transcriptome changes in aneuploid cells and Bafilomycin A1 (Baf A1)-treated RPE1 cells. Circle color intensity represents correlation coefficient strength (p < 0.05 shown). (**H**) Representative western blot of PARP1 protein levels in control versus Baf A1-treated hCECs. (**I**) PARP1 mRNA levels (Log2) in cells treated with or without proteasome inhibitors (MG132 or bortezomib). Unpaired t-test for RPE1 and HCT116. One-way ANOVA with Dunnett's test for HCT116. ns, p>0.05; *p<0.05; **p<0.01; ***p<0.001; ****p<0.0001.

**Figure S6. PARP1 suppression in aneuploid cells and its relationship with drug resistance**

(**A**) Nuclear CEBPB signal in hCECs with or without Baf A1 treatment (50 nM, 24h). Left: representative immunofluorescence images of CEBPB (green), Phalloidin (red), and DAPI (blue) staining in control (untreated) or BafA1 treated hCECs (scale bar, 10 μm). Right: quantification of nuclear CEBPB in hCECs with or without Baf A1 treatment. Mean ± SD, n>100 cells per condition. Welch's t-test. (**B**) Nuclear CEBPB signal in near-diploid and aneuploid hCECs expressing control (shNC) or *TFEB* (shTFEB) or *TFE3* (shTFE3) -targetting shRNAs. Left: representative immunofluorescence images of CEBPB (green), Phalloidin (red), and DAPI (blue) staining in aneuploid hCEC with *TFEB* or *TFE3* knockdown (scale bar, 10 μm). Right: quantification of nuclear CEBPB in near-diploid and aneuploid hCECs with *TFEB* or *TFE3* knockdown. Mean ± SD, n>200 cells per condition. Kruskal-Wallis one-way Anova with Dunn's test. (**C**) Viability after H2O2 treatment (40 μM) in hCEC clones with *TFEB* or *TFE3* knockdown. Mean ± SD from 9 fields. One-way Anova with Sidak's test. (**D**) *PARP1* RNA levels in TCGA COREAD tumors stratified by *CEBPB* copy number status defined by cBioPortal, excluding tumors with PARP1 copy alterations. Mean ± SD. One-way ANOVA with Dunnett's test. (**E**) ChIP-qPCR of CEBPB enrichment at the *PARP1* promoter. Top: Schematic of the qPCR amplicons. Bottom: ChIP-qPCR analysis in DMSO (diploid), Reversine (aneuploid), and Dexamethasone (CEBPB agonist) treated hCECs. Data show mean ± SEM of % input values normalized to negative control, n=3. Unpaired t-test. (**F**) *CEBPB* and *PARP1* RNA level correlation in malignant colorectal cancer cells determined by single-cell RNA-seq analysis. Pearson correlation coefficient and p-value are indicated. (**G**) *CEBPB* and *PARP1* RNA level correlation in TCGA COREAD tumors. Linear regression is included to fit the data. (**H**) Schematic diagram illustrating the major findings of this study. Top: aneuploidy leads to PARP1 suppression through lysosomal stress-mediated CEBPB activation. Bottom: consequences of PARP1 suppression in aneuploid cells, including inhibition of cell death (parthanatos) and DNA repair pathways, and promotion of metastasis. (**I**) Top panels: PARP1 copy number, mRNA levels, and protein levels in cancer cell lines with different degrees of aneuploidy (CCLE dataset). Bottom panels: cell resistance to the PARP1 inhibitor olaparib across three different drug screen datasets–PRISM, CTD2, and GDSC2–in cells with different degrees of aneuploidy. Higher Z-scores correspond to greater resistance. Data are shown as Z-scores with box plots indicating median, first and third quartiles, and whiskers extending to 1.5× interquartile range. Individual data points are shown. One-way ANOVA with Tukey's multiple comparison test. For clarity, all p-values are included in Table 6.

### Supplementary Tables

**Table 1**. SCNAs of hCECs with acute and chronic aneuploidy. Details of drugs and COREAD cell lines used in Figure 1.
**Table 2**. Differential gene expression analysis of hCECs with acute and chronic aneuploidy.
**Table 3**. Statistical analysis of γH2AX intensity in near-diploid and aneuploid hCEC clones, with or without PARP1 overexpression, before and after $H_2O_2$ exposure.
**Table 4**. Aneuploidy degree, PARP1 RNA level and survival of TCGA COREAD patients.
**Table 5**. CRISPR screen of PARP1 regulators and pathway analysis.
**Table 6**. Transcription factor enrichment analysis of genes upregulated in aneuploid or lysosome-dysfunction cells
**Table 7**. Details of the materials and all statistical analyses in this study. Table 7 contains all data and statistical analyses for each panel shown in the paper. Each sheet is labeled with the corresponding figure number and panel.
